# Data-driven classification of tissue water populations by massively multidimensional diffusion-relaxation correlation MRI

**DOI:** 10.1101/2025.03.28.645938

**Authors:** Omar Narvaez, Maxime Yon, Raimo A. Salo, Jenni Kyyriäinen, Melina Estela, Ekaterina Paasonen, Ville Leinonen, Juhana Hakumäki, Frederik Laun, Daniel Topgaard, Alejandra Sierra

## Abstract

Massively multidimensional diffusion-relaxation correlation MRI provides detailed information on tissue microstructure by analyzing water populations at a sub-voxel level. This method correlates frequency-dependent tensor-valued diffusion MRI with longitudinal and transverse relaxation rates, generating nonparametric **D**(*ω*)-*R*_1_-*R*_2_ distributions. Traditionally, **D**(*ω*)-*R*_1_-*R*_2_ distributions are separated using manual binning of the diffusivity and anisotropy space to differentiate white matter (WM), gray matter (GM), and free water (FW) in brain tissue. However, while effective, this approach oversimplifies complex tissue fractions and does not fully utilize all available diffusion-relaxation parameters. In this study, we implemented an unsupervised clustering approach to automatically classify WM, GM, and FW and explore additional water populations using all components in the **D**(*ω*)-*R*_1_-*R*_2_ distributions on *ex vivo* and *in vivo* rat brain, and *in vivo* human brain. Results showed that a basic separation of WM, GM, and FW is possible using unsupervised clustering even under different multidimensional diffusion-relaxation protocols of rat brain and human brain. Additionally, when there is high frequency-dependent diffusion range, it is possible to obtain a cluster characterized by restriction localized in specific high cell density regions such as the dentate gyrus and cerebellum of rat brain. These findings were compared with rat histological sections of myelin and Nissl stainings. We demonstrated that unsupervised clustering of diffusion-relaxation MRI data can reveal tissue complexity beyond traditional WM, GM, and FW segmentation in rat and human brain without parameter assumptions. The unsupervised cluster approach could be used in other body parts (*e*.*g*., prostate and breast cancer) without requiring pre-defined bin limits. Furthermore, the characterization of the clusters by diffusivities, anisotropy, and relaxation rates can provide a better understanding of the subtle changes in different cellular fractions in tissue-specific pathologies.

## 1 Introduction

Diffusion magnetic resonance imaging (dMRI) provides a non-invasive means to access microstructural information about brain tissue by measuring the displacement of water molecules. The interaction of these water molecules with the complex cellular architecture and composition gives rise to multiple water populations associated with various cellular structures, including cell somas, neurites, and diverse lipidic and chemical compositions. The MRI signal from a single voxel reflects a distinct degree of heterogeneity, resulting from a sum of the contributions from these water populations. In pathological conditions, alterations in cellular structures—both in morphology and composition—can further increase the complexity of the dMRI signal by affecting the underlying water populations.

Pierpaoli and collaborators (Pierpaoli *et al*., 1996) in 1996 initially demonstrated the separation of meaningful water populations using only diffusion MRI data through diffusion tensor imaging parameters, trace and anisotropy. The use of per-voxel single trace and anisotropy values allowed to produce the segmentation of WM, GM, and FW regions. Expanding on this basis, the diffusion MRI community has made significant efforts in addressing intravoxel heterogeneity by developing both biophysical models (Novikov, Kiselev, *et al*., 2018; Jelescu *et al*., 2020) and advanced dMRI sequences (Lundell *et al*., 2019; Aggarwal, 2020; Reymbaut, 2020; Henriques *et al*., 2021; Jiang *et al*., 2023). Biophysical modeling separates water populations by defining cellular compartments. In white matter (WM), axons are often represented as sticks, extra-axonal space as an axially symmetric tensor, and free water (FW) as a spherical tensor (Zhang *et al*., 2012; Novikov, Veraart, *et al*., 2018; Coelho *et al*., 2022). In gray matter (GM), additional parameters account for the soma compartment (Palombo *et al*., 2020). Each model has specific assumptions, requiring compatible acquisition protocols for optimal application. Consequently, these models are limited to the water populations they are designed to analyze and may not be applicable to tissues outside the brain. In practice, studies focusing on WM require specific parameter estimators to capture different diffusion regimes, enhancing the fit between the models and the tissue signal. In addition to biophysical modeling, valuable information regarding water populations can also be obtained by adjusting the scanning acquisition parameters.

Recently, there has been an increased application of multidimensional acquisitions that combine b-tensor encoding diffusion and relaxation (De Almeida Martins *et al*., 2018, 2021; Martin *et al*., 2021). One approach is massively multidimensional diffusion-correlation MRI (Narvaez *et al*., 2022), which facilitates the acquisition of per-voxel nonparametric distributions of diffusivities, anisotropies, restrictions, and chemical compositions (**D**(*ω*)-*R*_1_-*R*_2_-distributions). In this approach, the signal is obtained using free gradient waveforms by modifying the trace (*b*), normalized anisotropy (*b*_Δ_), orientation (Θ, Φ), centroid frequency (*ω*_cent_) with variable repetition (τ_R_), and echo time (τ_E_). Once the distributions are obtained, an arbitrary set of “bins” limits are selected on the isotropic diffusivity (*D*_iso_) and squared normalized anisotropy (*D*_Δ_^2^) space to separate and visualize water populations into meaningful tissue fractions (*i*.*e*., WM and GM) and free water (FW). Similarly, earlier binning approaches included 1D R_2_-distributions for separating myelin water fractions (Kim *et al*., 2017) or 2D “spectral regions of interest” for differentiating tissue types from 2D correlation experiments (Benjamini *et al*., 2017; Slator *et al*., 2019; Pas *et al*., 2020). In nonparametric **D**(*ω*)-*R*_1_-*R*_2_-distributions, *D*_iso_ and *D*_Δ_^2^ contains homologous microstructural information to mean diffusivity (MD) (Basser *et al*., 1994). and microscopic fractional anisotropy (µFA) (Lasič *et al*., 2014; Shemesh *et al*., 2016a; Topgaard, 2016). Even though the “bin” division of the per-voxel distributions is practical and yields adequate separation of WM-like, GM-like and FW-like water populations, the selection of the “bin” limits needs to be manually modified in order to provide meaningful tissue fractions under different conditions (i.e., *in vivo, ex vivo*, type of sample analyzed); and it misses the opportunity to exploit the multiple dimensions available. Moreover, the “bin” selection falls short in disentangling the complexity inherent within the voxel. In addition to well defined water population in tissue, there are potential image artifacts and not physical plausible distributions due to under-sampling of *b*_Δ_ at low *b*-values (De Almeida Martins *et al*., 2018) that could be mixed with true water population using manual binning approaches. This limitation highlights the need for unsupervised methods capable of capturing biological relevant features from multidimensional diffusion-relaxation correlation experiments.

Previous studies have utilized unsupervised approaches incorporating full tensor distribution data to further characterize tissue heterogeneity in diffuse gliomas (Song *et al*., 2022). Full tensor distributions account for both the symmetric and asymmetric components of the tensor, in contrast to diffusion tensor distribution (Topgaard, 2019) (DTD) approaches, which consider only the axially symmetric components. Most recently, the simultaneous acquisition of diffusion data while modifying T1 and T2 relaxation times has demonstrated the ability to differentiate water populations and provide greater specificity in the context of astrogliosis (Benjamini *et al*., 2023). Kundu and collaborators (Kundu *et al*., 2023) have introduced an unsupervised learning approach on the optimal transpose distances (LOT) from the multidimensional T1-T2, T1-MD and T2-MD spaces to distinguish cortical layers that could not be distinguishable using only 1D relaxation or diffusion parameters.

In this study, we used Gaussian mixture model (GMM) to perform data-driven clustering of nonparametric multidimensional diffusion-relaxation distributions, extending beyond the traditional fractions of WM, GM, and FW. We explore the classification of WM, GM, and FW tissue fractions while also identifying additional detailed tissue clusters within both WM and GM. These additional clusters can represent shared region-specific water diffusion populations or possible artifacts within a voxel. The data-driven clustering was applied to a comprehensive acquisition protocol for *ex vivo* rat brain data, as well as shorter protocols for both *in vivo* rat and human brains. This work represents a significant advancement in understanding the efficacy of this approach compared to previous methodologies.

## 2 Materials and methods

### 2.1 Animals and data acquisition

#### 2.1.1 *Ex vivo* healthy rat brain at 11.7-T scanner

An adult male Sprague-Dawley rat was used for the *ex vivo* study (10 weeks old, 350 g, Harlan Netherlands B.V.). The rat was housed individually in a cage with climate-control room under a 12h/12h light/dark cycle with *ad libitum* diet. The animal procedures were approved by the Animal Ethics Committee of the Province Government of Southern Finland and carried out according to the guidelines set by the European Community Council Directives 2010/63/EEC.

First, the rat was anesthetized with 5% isoflurane in a mixture of 70% nitrogen/30 % oxygen and transcardially perfused with 0.9% NaCl for 5 min (30 ml/min) followed by 4% paraformaldehyde (PFA) in 0.1 M phosphate buffer (PB) pH 7.4 for 25 min (30 ml/min). After the perfusion, the brain was removed from the skull and postfixed in 4% PFA for 4 h. Before the *ex vivo* imaging, the brain was immersed in 0.1 M phosphate buffer saline (PBS) pH 7.4 to remove the excess of PFA for 24 h. Only before the scan, the brain was divided into two hemispheres. Then, the left hemisphere was placed inside a 10-mm diameter tube filled with perfluoropolyether (Galden, Solvay, Italy) to suppress background signal in the images.

The dataset was acquired on a Bruker Avance-III HD 11.7-T spectrometer with a MIC-5 probe giving 3 T/m maximum gradient amplitude on-axis using a 10-mm diameter coil. The images were obtained using Bruker’s multi-slice multi-echo (MSME) sequence customized to use double-rotation gradient waveforms (Jiang *et al*., 2023) running on Paravision 6.01. The multidimensional acquisition of the rat brain consisted of 1,095 images varying *b*-value (0.038-7.13·10^9^ sm^−2^), centroid frequency *ω*_cent_/2π within 10-90% (29-110 Hz), normalized anisotropy *b*_Δ_ (–0.5, 0, 0.5, and 1), orientation (Θ,Φ), recovery time τ_R_ (200-7000 ms), and echo time τ_E_ (6-180 ms) in a total scanning time of 87 h. The whole acquisition protocol is shown in **Supplementary Figure 1A**. The images were acquired at 18 ± 1 °C. The acquisition was a single slice in the coronal plane with a resolution of 70 × 70 × 250 µm^3^ (matrix size 122 × 128 × 1) located at approximately –3.5 mm from bregma.

The acquired images were preprocessed as follows: The data was reconstructed using Paravision 6.01 into the real and imaginary domains and denoised, using MPPCA approach included in DESIGNER toolbox (Chen *et al*., 2024). After denoising, the magnitude image was calculated and Gibbs-ringing artifact removal was applied with the sub-voxel shifts (Kellner *et al*., 2016) using MRtrix3 toolbox (Tournier *et al*., 2019).

#### 2.1.2 *In vivo* rat brain with at 7-T scanner

One male Sprague-Dawley rat was used for the *in vivo* experiment (10 weeks old, 350 g, Harlan Netherlands B.V). It was housed individually in a cage with climate-control room under a 12h/12h light/dark cycle with *ad libitum* diet. The animal procedures were approved by the Animal Ethics Committee of the Province Government of Southern Finland and carried out according to the guidelines set by the European Community Council Directives 2010/63/EEC.

The rat was anesthetized under ∼5% isoflurane in a mixture of 70% nitrogen/30% oxygen for 5 min, after which it was placed on the animal bed equipped with heated water circulation. The anesthesia then was maintained at ∼2% isoflurane with the same nitrogen/oxygen mixture. The body temperature (36-37°C) and respiration rate (35–60 breaths per minute) were monitored during the scans.

The dataset was acquired on a 7-T/16-cm horizontal Bruker PharmaScan 7 Tesla preclinical system with a maximum gradient strength of 760 mT/m running on Paravision 6.01. An actively decoupled standard volume coil was used for RF-transmission and a quadrature surface coil for receiving. The images were obtained using Bruker’s Spin Echo-Echo Planar Imaging (SE-EPI) sequence customized to use double-rotation gradient waveforms (Jiang *et al*., 2023; Yon *et al*., 2025). The images were acquired in a horizontal plane with a resolution of 250 × 250 × 700 µm^3^ with a slice gap of 300 µm and matrix size of 80 × 129 × 5. Double sampling and two segments were used. The images were acquired with two repetitions to include the reversed phase-encode images for later preprocessing steps. The multidimensional acquisition of the rat brain consisted of 389 pair of images varying *b*-value (0.031-2.65·10^9^ sm^−2^), centroid frequency *ω*_cent_/2π within 10-90% (18-92 Hz), normalized anisotropy *b*_Δ_ (–0.5, 0, 0.5, and 1), orientation (Θ,Φ), repetition time τ_R_ (1000-3500 ms), and echo time τ_E_ (21-77 ms) in a total scanning time of 1 h 9 min. The whole acquisition protocol is shown in **Supplementary Figure 1B**.

The preprocessing of the *in vivo* rat images was as it follows: 1) The images were reconstructed in Paravision 6.01. 2) The images were separated into their respective phase-encoding directions using Matlab (Natick, Massachusetts: The MathWorks Inc.) ; 3) each set of phase-encoding directions were denoised separately using the MPPCA approach (Chen *et al*., 2024); 4) Gibbs rings artifact removal (Kellner *et al*., 2016; Tournier *et al*., 2019); and 5) correction of susceptibility-induced off-resonance field using FSL top up tool as described in (Yon *et al*., 2025).

#### 2.1.3 *In vivo* human brain at 3-T scanner

Six healthy volunteers were scanned (2 females, 4 males: age range 23-51 years) in compliance with the Finnish ethical permit 134/2023, 23.05.2023 and Finnish Medicines Agency (FIMEA) permit, FIMEA/2023/004206 in a 3-T MAGNETOM Vida Siemens scanner, equipped with a 70 cm bore size and 60/200 XT gradients. T1w images with voxel size of 1 × 1 × 1 mm^3^, repetition time τ_R_ = 2.3 s, and echo time τ_E_ = 1.9 ms were evaluated by an experienced radiologist, and no abnormalities were detected. The multidimensional images were obtained with a SE-EPI sequence customized to use diffusion general gradient waveforms(Martin *et al*., 2020). The data was acquired with a voxel size of 2 × 2 × 2 mm^3^ (matrix size 114 × 114 × 55 mm^3^). The multidimensional acquisition of the human brain consisted of 139 images varying *b*-value (0-3·10^9^ sm^−2^), centroid frequency *ω*_cent_/2π within 10-90% (4-9 Hz), normalized anisotropy *b*_Δ_ (–0.5, 0, and 1) using numerically optimized waveforms (Sjölund *et al*., 2015), orientation (Θ, Φ), repetition time τ_R_ (620-7000 ms), and echo time τ_E_ (40-150 ms) in a total scanning time of 40 min. The whole acquisition protocol is shown in **Supplementary Figure 1C**.

The preprocessing of the *in vivo* human images was as it follows: 1) The images were denoised using the MPPCA approach (Chen *et al*., 2024); 2) Gibbs rings artifact removal (Kellner *et al*., 2016; Tournier *et al*., 2019); and 3) correction of susceptibility-induced off-resonance field using FSL top up tool using a couple of non-diffusion weighted images with two different phase-encoding directions (Andersson *et al*., 2003).

### 2.2 Estimation of nonparametric D(*ω*)-*R*_**1**_**-*R***_**2**_-distributions and visualization

Nonparametric **D**(*ω*)-*R*_1_-*R*_2_-distributions are derived through the approximation of the **b**(*ω*)-τ_R_-τ_E_ encoded signal *S* according to the following equation:

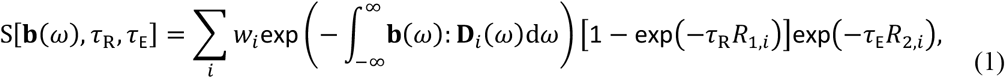

The inversion of **Eq. 1** allows the acquisition of the ω-dependent self-diffusion tensor **D**(*ω*) by approximating **D**_*i*_(*ω*) as an axisymmetric tensor,

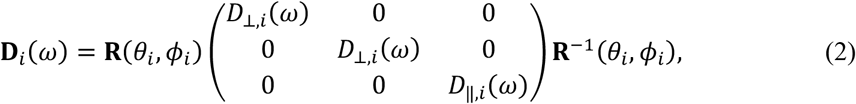

where **R**(*θ*_*i*_, *ϕ*_*i*_) is a rotation matrix and ω-dependent parallel and perpendicular eigenvalues, *D*_‖,*i*_(*ω*) and *D*_⊥,*i*_(*ω*), are described by Lorentzian transitions (Narvaez *et al*., 2024) according to:

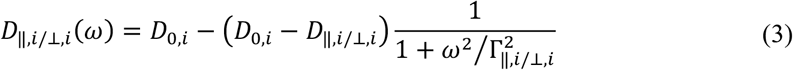

Using the Lorentzian approximation, each component in the **D**(*ω*)-*R*_1_-*R*_2_-distributions is described with its weight *w* and parameter set [*D*_∥_, *D*_⊥_, *θ, ϕ, D*_0_, Γ_∥_, Γ_⊥_, *R*_1_, *R*_2_]. We use the Monte Carlo inversion (Prange *et al*., 2009) described in detail in (Narvaez *et al*., 2022) and (Narvaez *et al*., 2024) to retrieve the ensemble of solutions using Matlab (Natick, Massachusetts: The MathWorks Inc.) with the *md-dmri* toolbox (Nilsson *et al*., 2018).

The inversion limits used for the *ex vivo* rat brain at 11.7 T, *in vivo* rat at 7 T, and *in vivo* human brain at 3 T are presented in **Table 1**.

**Table 1.**
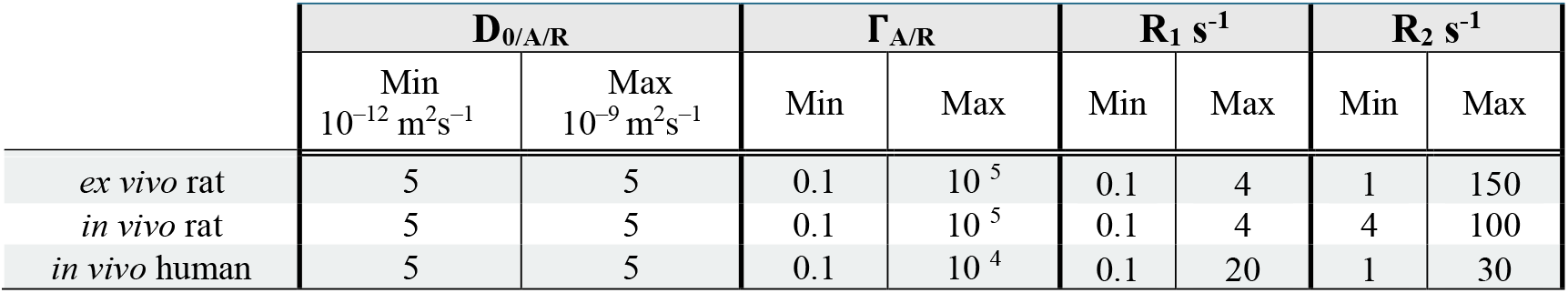
Monte Carlo inversion limits.

The Monte Carlo inversion settings for the three experiments were set to 20 steps of proliferation, 20 steps of mutation/extinction, 200 input components per step of proliferation and mutation/extinction, 10 output components, and bootstrapping by 100 repetitions using random sampling with replacement. As a result, the Monte Carlo inversion outputs an ensemble of 100 independent per-voxel solutions. Each of the solutions comprised the weights *w* and coordinates [*D*_∥_, *D*_⊥_, *θ, ϕ, D*_0_, Γ_∥_, Γ_⊥_, *R*_1_, *R*_2_] for 10 components within the pseudo-randomly sampled primary analysis space. We then evaluated **D**(*ω*) distributions at chosen values within 10 and 90 percentiles of the *ω* range probed by the gradient waveforms (Narvaez *et al*., 2022, 2024; Yon *et al*., 2025), giving *ω*-dependent distributions [*D*_∥_(*ω*), *D*_⊥_(*ω*), *θ, ϕ, R*_1_, *R*_2_]. To follow well stablished metrics in the field (Pierpaoli *et al*., 1996), we obtained isotropic diffusivity *D*_iso_(*ω*), normalized anisotropy *D*_Δ_(*ω*), and squared normalized anisotropy *D*_Δ_^2^(*ω*) through:

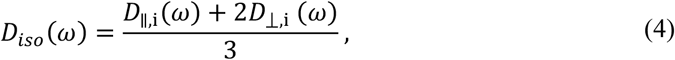

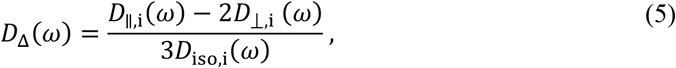

and

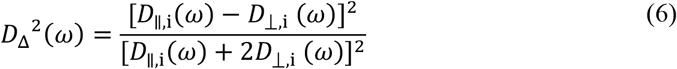

respectively.

In line with previous results from oscillating gradient encoding (Aggarwal *et al*., 2012, 2020; Xu *et al*., 2016; Aggarwal, 2020; Arbabi *et al*., 2020) and frequency-dependent multidimensional studies (Narvaez *et al*., 2022, 2024; Johnson *et al*., 2024; Manninen *et al*., 2024; Yon *et al*., 2025), restriction effect in *ω*-dependent *D*_iso_ and *D*_Δ_^2^ is quantified using the difference within the investigated frequency window from low to high *ω*.

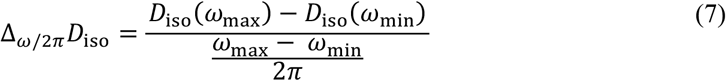

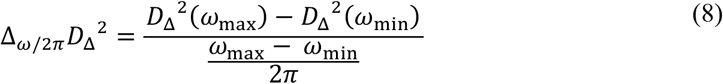

The distributions can be visualized using contour plots in the 2D *D*_*iso*_-*D*_Δ_^2^, *D*_*iso*_-*R*_1_, and *D*_*iso*_-*R*_2_ space, as well as bin-resolved signal fractions *f*_bin*n*_, means E_bin*n*_ [*X*], variances V_bin*n*_ [*X*], and covariances C_bin*n*_ [*X,Y*] appropriate to create per-voxel parameter maps (Topgaard, 2019). In this study, we used only per-voxel *f*_bin*n*_ and E_bin*n*_ [*X*] for visualization, calculated as follows:

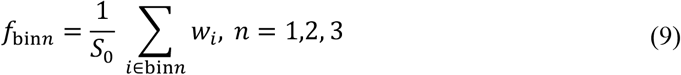

and

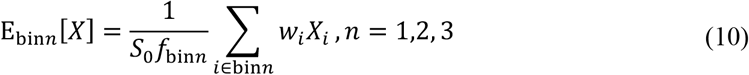

### 2.3 Unsupervised clustering of D(*ω*)-*R*_**1**_**-*R***_**2**_-distributions

Derived from 100 bootstraps with 10 components each, we obtain a total of 1000 per-voxel components. To consider the weights (*w*), each parameter value (*D*_iso_, *D*_Δ_ or *D*_Δ_^2^, *R*_1_, *R*_2_, and Δ_ω/2π_*D*_iso_) was replicated by its associated weight normalized and discretized between 0 and 100. Then, the resulted weighted parameters *D*_iso_, *D*_Δ_ or *D*_Δ_^2^, *R*_1_, *R*_2_, and Δ_ω/2π_*D*_>iso_ were organized into a 2D matrix and clustered using GMM approach, which is a probabilistic model that assumes that the data is a result of a mixture of normally distributed processes.

Using multidimensional normal distribution,

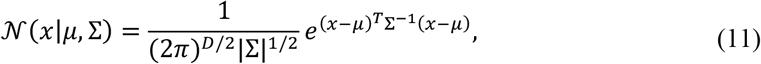

where *µ* is the mean of the distribution, Σ is the standard deviation of the distribution, and *x* is data, the GMM can be written as

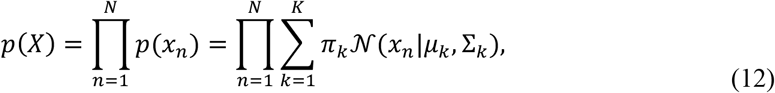

where *x*_*n*_ is a data sample, *K* is the number of clusters, *N* is the number of datapoints, and ∑_*k*_ π_*k*_ = 1. The GMM was solved by an expectation maximization algorithm that maximizes the likelihood of a model given the number of mixture components *k*. The different dimensions in the bootstrapped data were normalized to the same range by dividing each variable value with the standard deviation of the variable.

The data was clustered in two different ways. In the first one, we only include the variables *D*_iso_ and *D*_Δ_^2^ into the clustering using *k* = 3 to compare the results with the manual “3-bin” resolved fractions as used in previous studies (De Almeida Martins *et al*., 2020; Yon *et al*., 2020). For the second approach, we used *D*_iso_, *D*_Δ_, *R*_1_, *R*_2_ and Δ_ω/2π_*D*_iso_ variables for the *ex vivo* and *in vivo* rat brain, and *D*_iso_, *D*_Δ_, *R*_1_, *R*_2_ for the *in vivo* human brain using *k =* 2-10. We selected *D*_Δ_ for clustering to clearly differentiate the oblate distribution, facilitating the possible identification of inversion artifacts or biologically relevant features. As it is well known, unsupervised clustering methods rely in the selection of the *k* components. We calculated Schwarz’s Bayesian Information Criterion (BIC) to aid the selection of the ideal *k* components using *k =* 2-15 **Supplementary Figure 2**. The spatial information from the resulting clusters, clus*n*, can be visualized using the per-voxel normalized frequency, Fclus*n*. Using a similar formulation as bin-resolved approach, we obtained the cluster-resolved signal fractions *f*_clus_ which can be visualized using contour plots in the 2D *D*_iso_-*D*_Δ_^2^, *D*_iso_-*R*_1_, and *D*_iso_-*R*_2_ space and per-voxel means E_clus*n*_ [*X*] parametric maps calculated as follow:

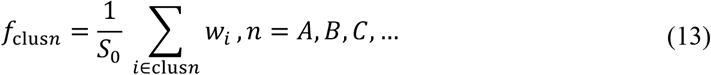

and

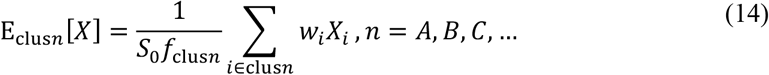

In the *in vivo* human data, we computed the clusters for every subject independently to compare the cluster centroids and covariance matrices across solutions.

### 2.4 Tissue processing and histology

The hemisphere scanned *ex vivo* was washed 2 times for 25 min in 0.1 M PB pH 7.4 and then cryoprotected for 48 h in a solution containing 20% glycerol in 0.02 M potassium phosphate-buffered saline (KPBS) pH 7.4. After cryoprotection, the hemisphere was frozen in dry ice and stored at –70°C until further processing. The hemisphere was coronally sectioned into 30-µm thick section and 1:2 series using sliding microtome. The first series of sections were stored in 10% formalin at room temperature and used for Nissl staining using thionin. The second series of sections were collected into tissue collection solution (30% ethylene glycerol, 25% glycerol in 0.05 M sodium phosphate buffer) and stored at –20°C until staining with gold chloride for myelin (Laitinen *et al*., 2010).

### 2.5 MRI region-of-interest (ROI) and histology analyses

ROI analysis was performed on the *ex vivo* rat brain using the cluster fractions resulting from the full space clustering with *k* = 7. ROIs were selected in the auditory cortex, primary somatosensory cortex, granule cell layer of the dentate gyrus, corpus callosum and cingulum. *D*_iso_, *D*_Δ_, *R*_1_, and *R*_2_ distributions of *f*_clusA_, *f*_clusB_, *f*_clusC_, *f*_clusD_, and *f*_clusG_ were obtained for each ROI. The projections were normalized by the maximum intensity within the cluster fractions. This normalization allowed to compare the cluster fraction contribution to each ROI independently of the ROI size.

We used a Nissl-stained section and a myelin-stained section from the same level in MRI to perform a comparison of cellular features with ROIs. Cells segmentation was performed on the Nissl-stained section, in which the per-cell area was calculated using QuPath v.0.5.0 software (Bankhead *et al*., 2017). Structure tensor (ST) analysis was calculated on the myelin staining (Budde *et al*., 2012; Molina *et al*., 2020), then, the photomicrograph was divided in patches of 70 × 70 μm^2^ to match with the in-plane voxel resolution as in *ex vivo* MRI. Angular central Gaussian (ACG) probability method was used to parametrize the results derived from the ST (Salo *et al*., 2021) and the coherency map 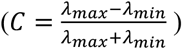 was calculated (Püspöki *et al*., 2016). In addition, the optical density (OD) on the myelin staining was computed by manually selecting a background threshold and normalizing.

## 3 Results

**Figure 1A and B** shows the division of **D**(*ω*)-*R*_1_-*R*_2_ distributions into the manual three “bin” limits on the *D*_iso_-*D*_Δ_^2^ space, as in previous studies (De Almeida Martins *et al*., 2018, 2020, 2021; Yon *et al*., 2020; Martin *et al*., 2021; Reymbaut *et al*., 2021; Narvaez *et al*., 2022, 2024). These bins can be considered as WM-like (bin1), GM-like (bin2) and FW-like (bin3). In the present study, the term WM to refer to such as the corpus callosum, cingulum, internal capsule, and optic tract (see the myelin-stained section in **Figure 6A and C**). Bin1 also included highly myelinated regions in the deep cortical and thalamic areas. Bin2 represented the remaining GM regions of the brain, while bin3 included the ventricles and liquid on the periphery of the brain.

**Figure 1.**
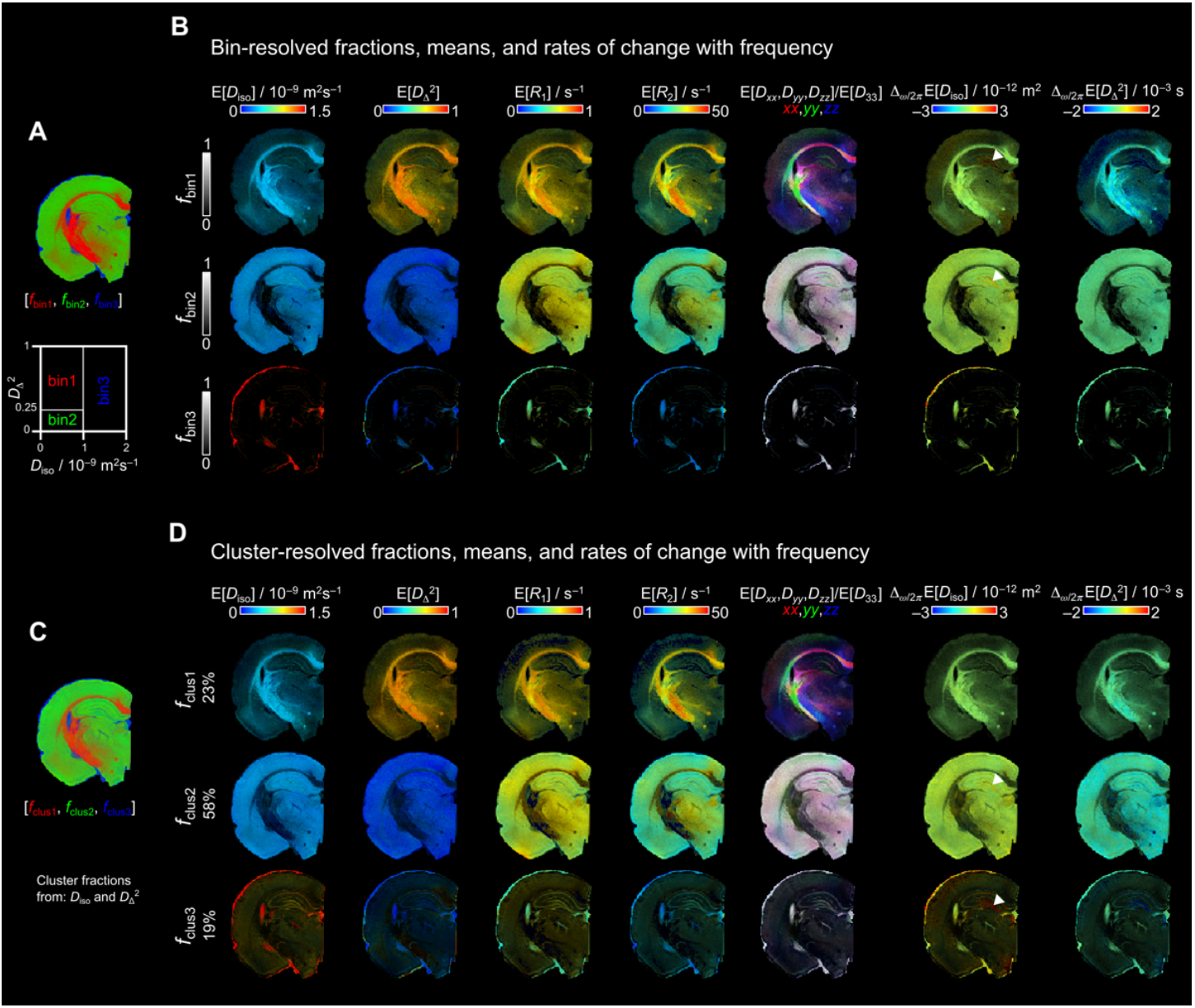
Bin- and cluster-resolved parameter maps derived from D(*ω*)-*R*_1_-*R*_2_ distributions. **(A)** Bin fractions segmenting WM (red), GM (green) and FW (blue) regions by manually setting the “bin” limits in the *D*_iso_-*D*_Δ_^2^ space. **(B)** Bin-resolved parametric maps for each “bin” fraction. The brightness represents the signal fractions, and the color scales the values of each parameter. (**C**) Color-coded unsupervised cluster-resolved normalized fractions by setting *k =* 3 using the *D*_*i*so_ and *D*_Δ_^2^ distributions. WM, GM, and FW fractions are displayed in same colors as the bin fractions. **(D)** Cluster-resolved parametric maps for each cluster fraction. Each fraction shows in percentage the amount of data belonging to the respective cluster. Arrowheads in Δ_ω/2π_E[*D*_iso_] maps indicate the granule cell layer of the dentate gyrus. Both bin- and cluster-resolved maps were calculated using the **D**(*ω*)-*R*_1_-*R*_2_ distributions at a frequency *ω*/2π = 29 Hz.

Our first step was to compare the bin-resolved mean parametric maps, and the maps obtained with our unsupervised classification approach using *k* = 3. **Figure 1A and B** shows the per-voxel mean parametric maps of bin1 with characteristic low E[*D*_iso_], high E[*D*_Δ_^2^], and both high E[*R*_1_] and E[*R*_2_] values in WM. bin2 had similar E[*D*_iso_] as in bin1, and lower E[*D*_Δ_^2^], E[*R*_1_] and E[*R*_2_] than in bin1, as expect in GM. bin3 showed higher E[*D*_iso_] than in bin1 and bin2, while E[*D*_Δ_^2^], E[*R*_1_], and E[*R*_2_] values were lower than in bin1 and bin2, corresponding to FW. **Figure 1C and D** display the cluster-resolved mean parametric maps using *k* = 3 in the *D*_iso_-*D*_Δ_^2^ space. The parametric maps of three cluster-fractions resemble the bin-resolved counterparts. The major difference between these two approaches was found in the FW fraction (**Figure 1D**). bin3 was only present in the ventricles and brain periphery, while clus3 showed the ventricles, brain periphery but also GM. In clus3, the highest E[*D*_iso_] values were found in the ventricles and the periphery of the brain. In GM, the E[*D*_iso_] values were lower than in clus3, but not as low as in clus2. The frequency-dependent parameter, Δ_ω/2π_E[*D*_iso_], also showed differences between bin- and cluster-resolved maps. While the highest Δ_ω/2π_E[*D*_iso_] values were mainly found in the granule cell layer of the dentate gyrus in the bin-resolved bin1 and bin2, these high values were found in the cluster-resolved clus2 and clus3 (white arrowhead in **Figure 1**).

**Figure 2** displays the resulted normalized cluster frequency by setting *k* from 2 to 10 when applied the full *D*_iso_, *D*_Δ_, *R*_1_, *R*_2_ and Δ_ω/2π_*D*_iso_ space. As showed in **Figure 1D**, using the *D*_iso_-*D*_Δ_^2^ space generated biological relevant fractions (*i*.*e*., a clear distinction of WM, GM, and FW regions) using *k* = 3; however, when using the full space required at least to *k* = 4 to get such biologically relevant fractions.

**Figure 2.**
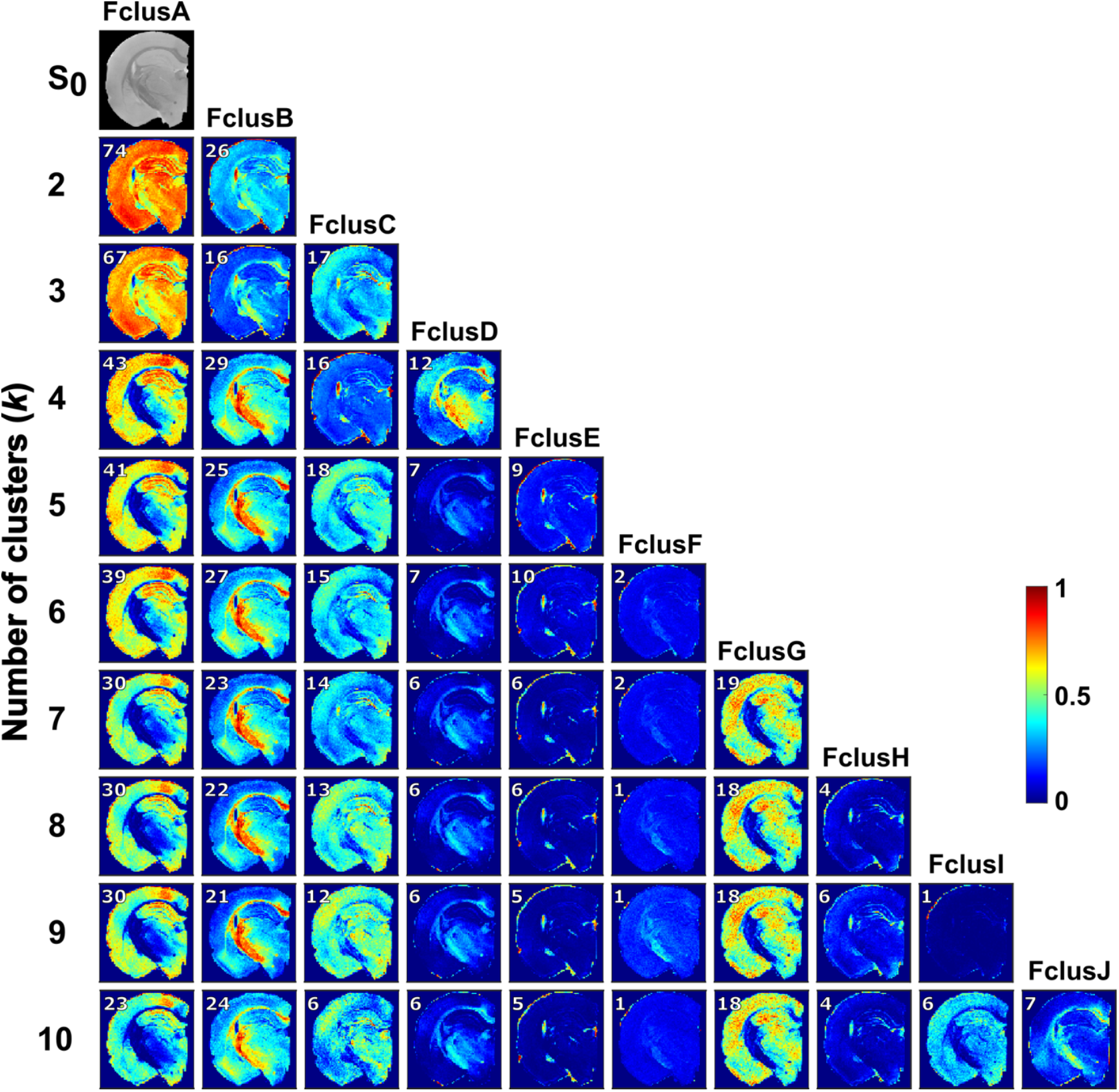
Normalized cluster frequency of *k* = 2-10 from the full distribution space (*D*_iso_, *D*_Δ_, *R*_1_, *R*_2_ and Δ_*ω*/2π_*D*_iso_). Per-voxel cluster frequency using *k* from 2 to 10 (rows show the number of clusters and columns the cluster fractions). The color scale shows the per-voxel belonging to each cluster.

Using *k* = 4, FclusA (included 43% of the data) contains mainly GM, while major WM regions were in FclusB (29%) including the corpus callosum, external and internal capsule or optic tract. FclusC (16%) comprised FW regions in the ventricles and periphery of the brain, and the granule and pyramidal cell layers. FclusD (12%) contained WM areas, especially in the cingulum and internal capsule, as well as highly myelinated thalamic regions. The difference between *k* = 4 and *k* = 5 clusters was that FclusC in *k* = 4 split into 2 fractions in *k* = 5: FclusC displaying GM regions and FclusE mainly ventricles and periphery of the brain. clusD in *k* = 5 exclusively showed WM regions. With *k* = 6, an additional cluster, FclusF (2%), was identified throughout the entire brain. Using *k* = 7, we found the separation of FclusA (30%) into FclusG (19%) displaying GM regions. The clusters obtained with *k* = 8, 9 and 10 remained constant as with *k* = 7, with the addition of FclusH, FclusI and FclusJ; however, these clusters contain small percentages of data of approximately 1-7%.

From *k* = 7 onwards, the clusters FclusA to FclusG remained similar across regions and percentages. For this reason, we selected the *k* = 7 cluster to visualize the parameter distributions that define each of the fractions, and obtained 2D contour plots between *D*_iso_-*D*_Δ_, *D*_iso_-*R*_1_, and *D*_iso_-*R*_2_ (**Figure 3**). For plotting the frequency dependance, we used the frequencies at 10, 50 and 90 % (*ω* /2π = 29, 69.5 and 110 Hz). The cluster fractions were ordered by amount of data classified into each fraction from largest to smallest.

*f*_clusA_, *f*_clusG_, and *f*_clusC_ contained GM regions; however, those fractions were defined by different parameters (**Figure 3A**). *f*_clusA_ mainly consisted of *D*_Δ_ values close to zero; *f*_clusG_ contained negative *D*_Δ_ with the mode (Mo) = -0.46, and *f*_clusC_ was characterized by variable *D*_Δ_ at different frequencies *ω*/2π Hz. *f*_clusB_ and *f*_clusD_ were non-zero in WM regions, the main distinction between these two fractions was that *R*_1_ (Mo = 0.65 s^-1^) and *R*_2_ (Mo = 24.38 s^-1^) values in *f*_clusB_ were lower than in *f*_clusD_. In *f*_clusE_, non-zero values were present in FW regions with low *R*_1_ and *R*_2_ values than previous cluster fractions. *D*_iso_ values across cluster fractions showed that *f*_clusA_, *f*_clusB_, and *f*_clusG_ had a similar *D*_iso_ distributions (Mo = 0.27×10^-9^ m^2^s^-1^) (**Figure 3B**). *f*_clusC_ showed higher *D*_iso_ mode values at 0.62×10^-9^ m^2^s^-1^ at *ω*/2π = 110 Hz. FW cluster (*f*_clusE_) presented values over 1×10^-9^ m^2^s^-1^, while *f*_clusD_ showed the lowest *D*_iso_ values (Mo = 0.12 ×10^-9^ m^2^s^-1^) than the rest of the fractions.

**Figure 3.**
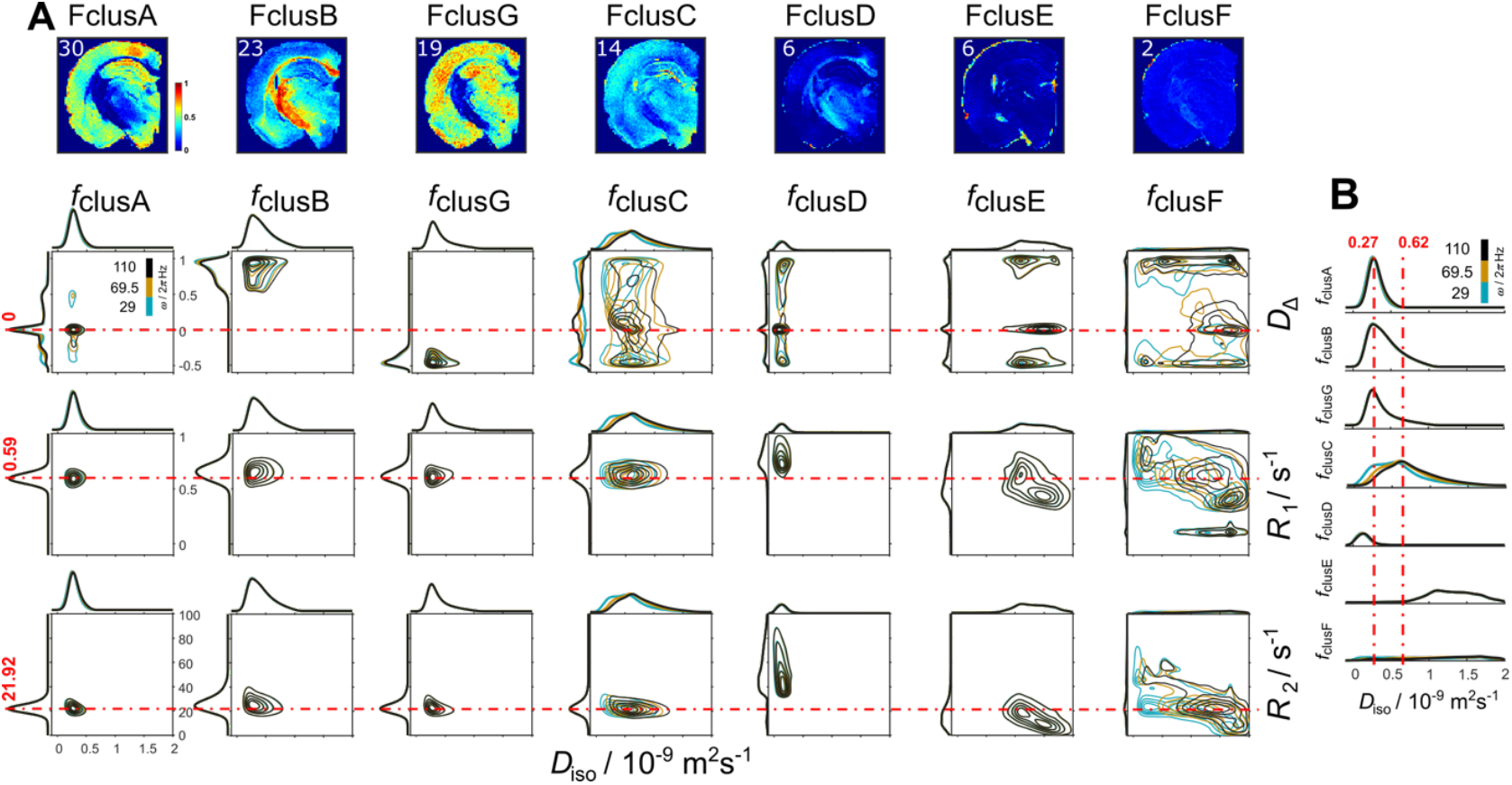
Per-cluster distributions for *k* = 7 derived from including *D*_iso_, *D*_Δ_, *R*_1_, *R*_2_ and Δ_*ω*/2π_*D*_iso_ into the clustering. **(A)** Per-voxel cluster frequency (columns) ordered from more percentage of data included into the cluster to the least percentage. The contour plots display the *D*_iso_-*D*_Δ_, *D*_iso_-*R*_1_, and *D*_iso_-*R*_2_ parameters (rows) for each cluster-fraction. Distributions at different ω/2π are plotted with different color (blue = 29, yellow = 69.5 and black = 110 Hz). The dashed red lines indicate the mode of the *D*_Δ_, *R*_1_, and *R*_2_ parameters from *f*_clusA_. **(B)** Distribution amplitude of *D*_iso_ dimension displays the comparison across cluster-fractions.

**Figure 4** shows all the parameter maps using the per-voxel cluster-resolved means in the *k* = 7 cluster fractions using the full space. We obtained the color-coded WM, GM, and FW-like fractions (**Figure 4A**) using the *f*_clusB_, *f*_clusA_ and *f*_clusE_, respectively. This visualization is a representation of the already described distributions in **Figure 3**. Hence, we can observe in **Figure 4B** WM-like regions (clusB and clusD) and GM-like regions (clusA, clusG and clusC) in color-coded according to the per-voxel means. It is worth emphasizing that previously, in **Figure 1B and D** the high Δ_ω/2π_E[*D*_iso_] were located mainly in the granule cell layer of the dentate gyrus within other tissue or FW fractions. Here (**Figure 4B**), clusC is the cluster that contains high Δ_ω/2π_E[*D*_iso_] not only in the granule cell layer but also across other GM-like regions.

**Figure 4.**
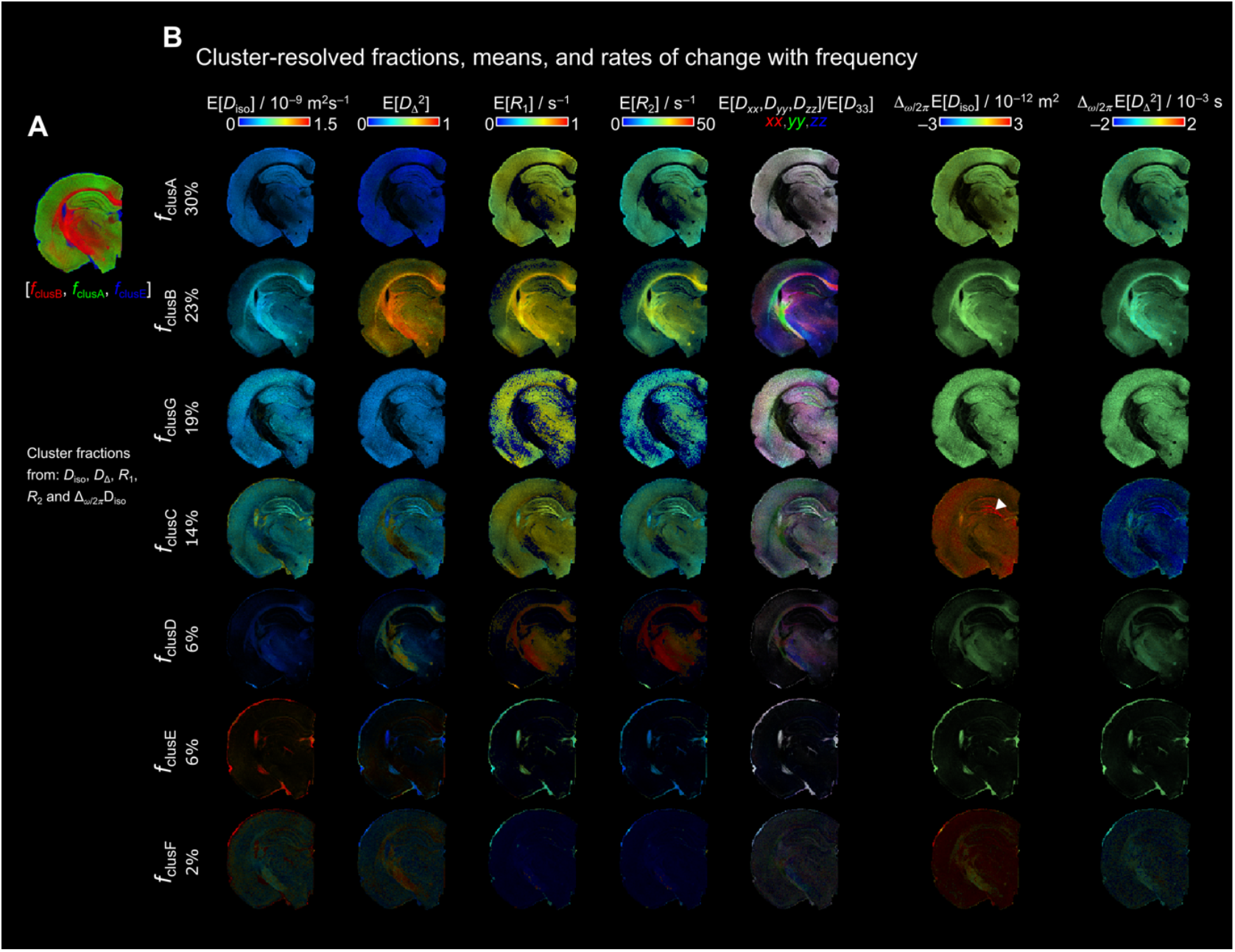
Cluster-resolved (*k* = 7) mean parameter maps derived from D(*ω*)-*R*_1_-*R*_2_ distributions. **A)** Unsupervised cluster-fractions *f*_clusB_, *f*_clusA_ and *f*_clusE_ in RGB color, respectively. This representation highlights the clusters-fractions that are WM, GM and FW-like. **(B)** Cluster-resolved parametric maps for each cluster-fraction. The cluster-fractions are ordered from higher to lower percentage of data belonging to each cluster. The color scales of the parameter maps are the same as in **Figure 1**. Arrowhead in Δ_ω/2π_E[*D*_iso_] map indicates the granule cell layer of the dentate gyrus. The cluster resolved maps were calculated using the **D**(*ω*)-*R*_1_-*R*_2_ distributions at a frequency *ω*/2π = 29 Hz.

The cluster-resolved parameter maps E[*D*_iso_], E[*D*_Δ_^2^], and E[*R*_2_] from the WM-like fractions clusB and clusD were compared with myelin staining from the same animal (**Figure 5A-C**). Although all are major WM fiber bundles, we observed differences in organization and density across the corpus callosum, cingulum, optic tract, and internal capsule. The optic tract demonstrated a more compact and dense organization than the corpus callosum. In contrast, the internal capsule exhibited complex crossing bundles, whereas to other regions like the corpus callosum were characterized by more uniformly parallel fiber organization (**Figure 5C**). The cingulum exhibited a rostro-caudal orientation, compared to the corpus callosum, which displayed an in-plane orientation in coronal sections (**Figure 5C**). Notably, although less dense than WM regions, certain GM regions—such as the thalamus, retrosplenial cortex, and deep cortical layers—exhibit high myelin content, as shown in **Figure 5A**. Regarding the parametric maps that can be comparable with the myelin staining by their presence in WM-like regions (**Figure 5B**), E[D_Δ_^2^] in clustB showed similar values across the corpus callosum, optic tract and internal capsule, while the E[*R*_2_] parametric map in clusD showed higher values in the optic tract and internal capsule than the corpus callosum (**Figure 5C**).

**Figure 5.**
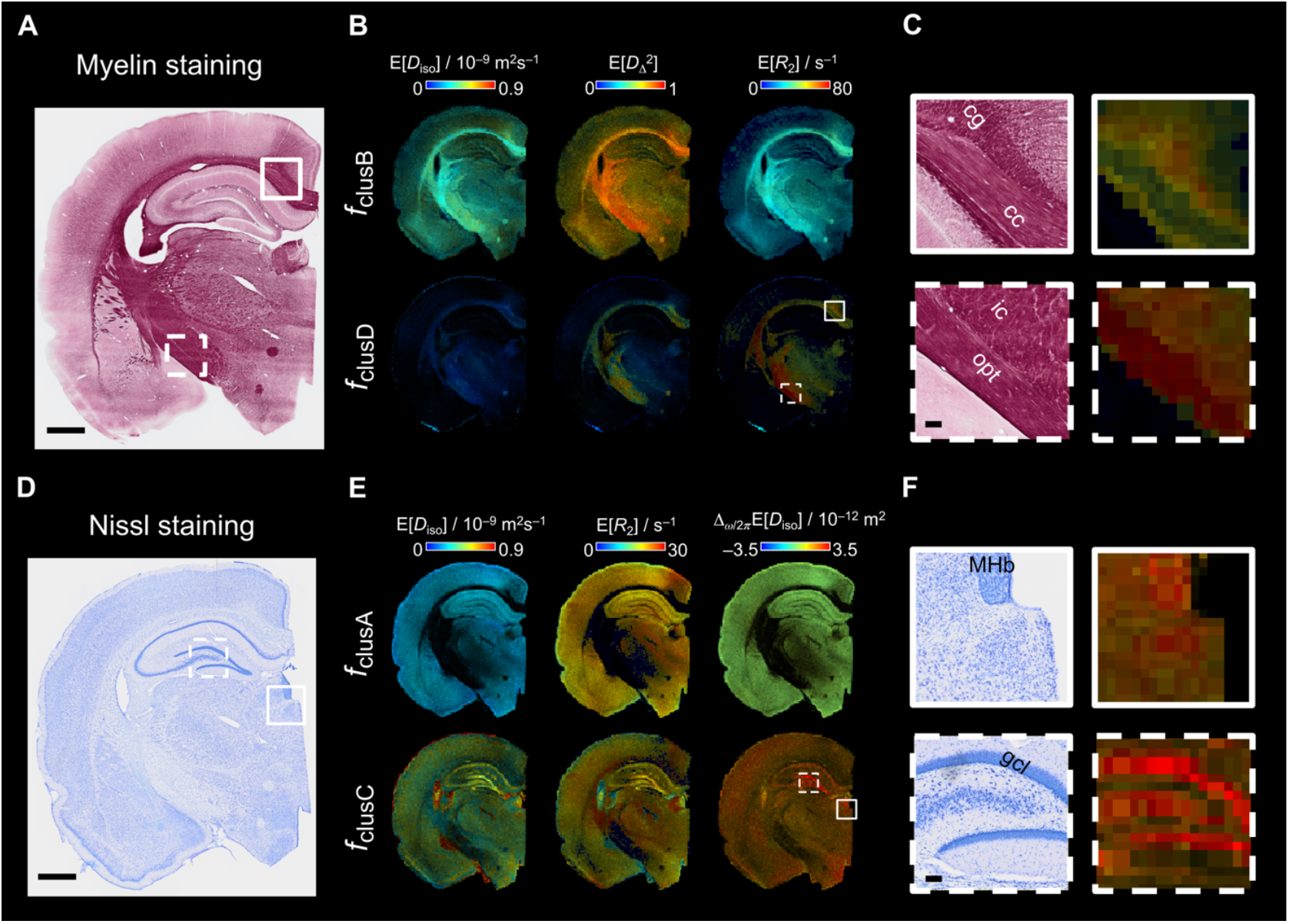
Histology and cluster-resolved mean maps comparison. **(A)** A representative myelin-stained section from the same brain as the acquired in MRI. Scale bar is 1 mm. **(B)** Representative cluster-resolved maps (E[*D*_iso_], E[*D*_Δ_^2^], and E[*R*_2_]) from **Figure 3**. The color scale bars limits are here modified from **Figure 3** to fit the scale values of the representative cluster fractions (*f*_clusB_ and *f*_clusD_). **(C)** Close up of two WM regions (continuous line square: corpus callosum (cc) and cingulum (cg); dashed line square: internal capsule (ic) and optic tract (opt)). Scale bar is 100 µm. **(D)** Representative Nissl-stained section from the same brain. Scale bar is 1 mm. **(E)** Representative cluster-resolved maps (E[*D*_iso_], E[*R*_2_], and Δ_ω/2π_E[*D*_iso_]). The color scale bars were modified as well from **Figure 3** to fit the scale values of the representative cluster fractions (*f*_clusA_ and *f*_clusC_). **(F)** Close up of two different GM regions (continuous line square: medial habenular nucleus (MHb); dashed line square: granule cell layer (gcl)). Scale bar is 100 µm.

The representative Nissl-stained section revealed regions with high cell density, such as the granule cell layer, CA3 and medial habenular nucleus (**Figure 5D and F**), which corresponded to high Δ_ω/2π_E[*D*_iso_] values in clusC (**Figure 5F**). In contrast, clusA exhibited limited differentiation between GM regions in the Δ_ω/2π_E[*D*_iso_] and E[D_iso_] parametric maps.

**Figure 6** presents the results of ROI analyses for selected brain areas using both MRI and histology, reveling distinct distribution patterns between WM and GM regions (**Figure 6B**). Here, the projections of the distributions were normalized by the maximum intensity within the cluster fractions. This normalization allowed to compare the proportion of data included in the cluster-fractions between ROIs independently of the number of voxels. *f*_clusA_ distributions for all four parameters (*D*_iso_, *D*_Δ_, *R*_1_ and *R*_2_) displayed consistent parameter values and similar data proportions across the auditory cortex, primary somatosensory cortex, and granule cell layer. The most notable difference emerged in *f*_clusC_, in which the proportion of data was larger in the granule cell layer, followed by the auditory cortex and then, by the primary somatosensory cortex. Comparison of GM regions with Nissl staining (**Figure 6C**) revealed the highest cell density in the granule cell layer. Although segmentation of individual cell bodies in the granule cell layer was not successful, the auditory cortex was found to have a larger cellular area compared to the primary somatosensory cortex (**Figure 6D**), consistent with observations in *f*_clusC_. Small differences were noted between GM regions in *f*_clusB_ and *f*_clusG,_ where the auditory cortex showed a higher data proportion than primary somatosensory cortex and granule cell layer. As mentioned above, *f*_clusB_ is associated with high myelin content. Within GM regions, the auditory cortex demonstrated higher density of myelinated axons, as showed by the myelin staining and optical density map (**Figure 6E and F**). The coherency map further revealed increased fiber orientation coherence toward the superficial cortical layers in both the auditory cortex and primary somatosensory cortex (**Figure 6G**).

**Figure 6.**
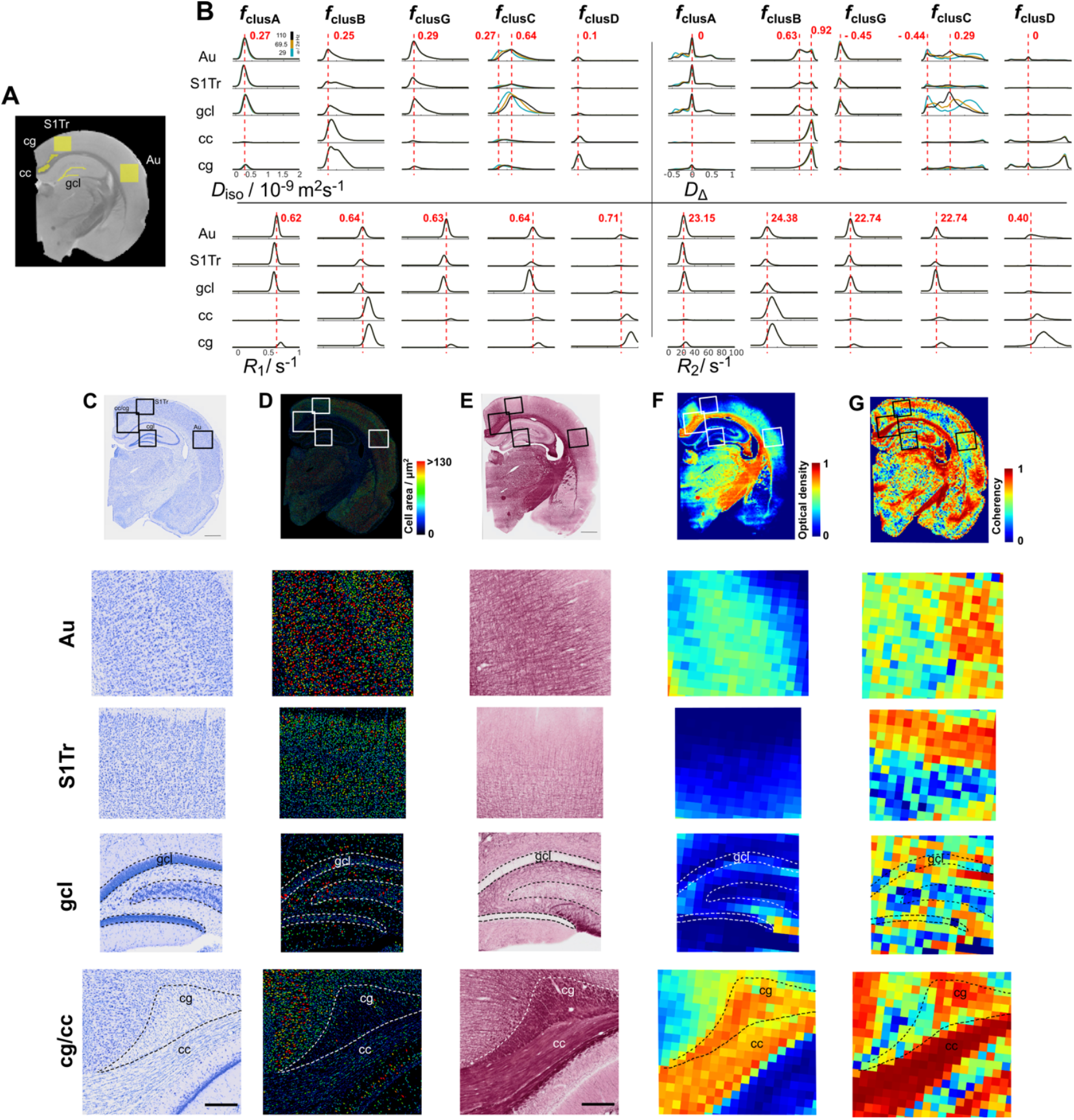
ROI analysis and comparison with Nissl and myelin staining. **(A)** selected ROIs overlayed on the S_0_ MRI. ROIs in GM: auditory cortex (Au), primary somatosensory cortex (S1Tr), and granule cell layer (gcl); ROIs in WM: corpus callosum (cc) and cingulum (cg). **(B)** per-ROI distributions for *f*_clusA_, *f*_clusB,_ *f*_clusG_, *f*_clusC_, and *f*_clusD_ ordered as in **Figure 3 and 4**. The per-ROI distributions are grouped by parameter. Upper left: *D*_iso_, upper right: *D*_Δ_, lower left: *R*_1_ and lower right: *R*_2_. Distributions at different ω/2π are plotted with different color (blue = 29, yellow = 69.5 and black = 110 Hz). Dashed red lines indicate the mode value in reference to the ROI in the auditory cortex. **(C)** Nissl-stained section from the same brain (scale bar is 1 mm) with close ups to the regions selected in the MRI slice (scale bar is 200 µm). **(D)** Cell area from the Nissl-stained section. **(E)** Myelin-stained section from the same brain (scale bar is 1 mm) with close ups to the regions selected in the MRI slice (scale bar is 200 µm). **(F)** Optical density map of the myelin-stained section. **(G)** Coherency map derived from the structure tensor analysis on the myelin-stained section. The analysis window was 70 × 70 µm^2^ to match the MRI voxel size.

The proportion of data within the corpus callosum and cingulum in *f*_clusA_, *f*_clusG_, and *f*_clusC_ was relatively small as compared to GM areas (**Figure6B**). However, among WM regions, the cingulum displayed a slightly higher proportion of data within those cluster-fractions than the corpus callosum. Both the corpus callosum and cingulum demonstrated lower cell density and smaller cellular area than GM regions (**Figure 6C and D**). In *f*_clusB_, the corpus callosum and cingulum showed consistent distributions across the *D*_Δ_, *R*_1_ and *R*_2_ parameters, with only small differences in *D*_iso_. Notably, the cingulum displayed a broader *D*_iso_ distribution compared to the corpus callosum. The most prominent difference was observed in *f*_clusD_, where the cingulum had a higher proportion of data and broader *R*_1_ and *R*_2_ distributions than the corpus callosum. Both, the corpus callosum and cingulum contained similar high densities of myelinated axons, as confirmed by the myelin staining and optical density map (**Figure 6E and F**). However, the coherency map revealed lower coherence values in the cingulum compared to the corpus callosum (**Figure 6G**).

We obtained the per-voxel cluster frequencies from the *in vivo* rat brain using *k* = 2 to *k* =10 (**Supplementary Figure 3**). Compared to the *ex vivo* rat brain data, the *in vivo* data were characterized by lower signal-to-noise ratio (SNR), larger voxel size, and a different acquisition protocol. Consequently, the optimal *k* value for biologically relevant clustering differed between *in vivo* and *ex vivo* datasets. Despite these differences, the resulting cluster at *k* = 6 (**Figure 7**) remained similar in terms of the overall parameter distributions.

In the *in vivo* dataset (**Figure 7A**), components belonging to clusA, clusB and clusC were primarily located in GM with smaller proportions in WM regions (33, 19 and 18%, respectively). clusD was observed in the ventricles and periphery of the brain (14%) while clusF was predominantly located in WM regions (13%). clusE was distributed across the entire brain but accounted for only a small portion of the clustered data (3%), consistent with findings from the *ex vivo* results that clusD of the *ex vivo* rat brain (**Fig.3A**) was not present in the *in vivo* clusters.

**Figure 7.**
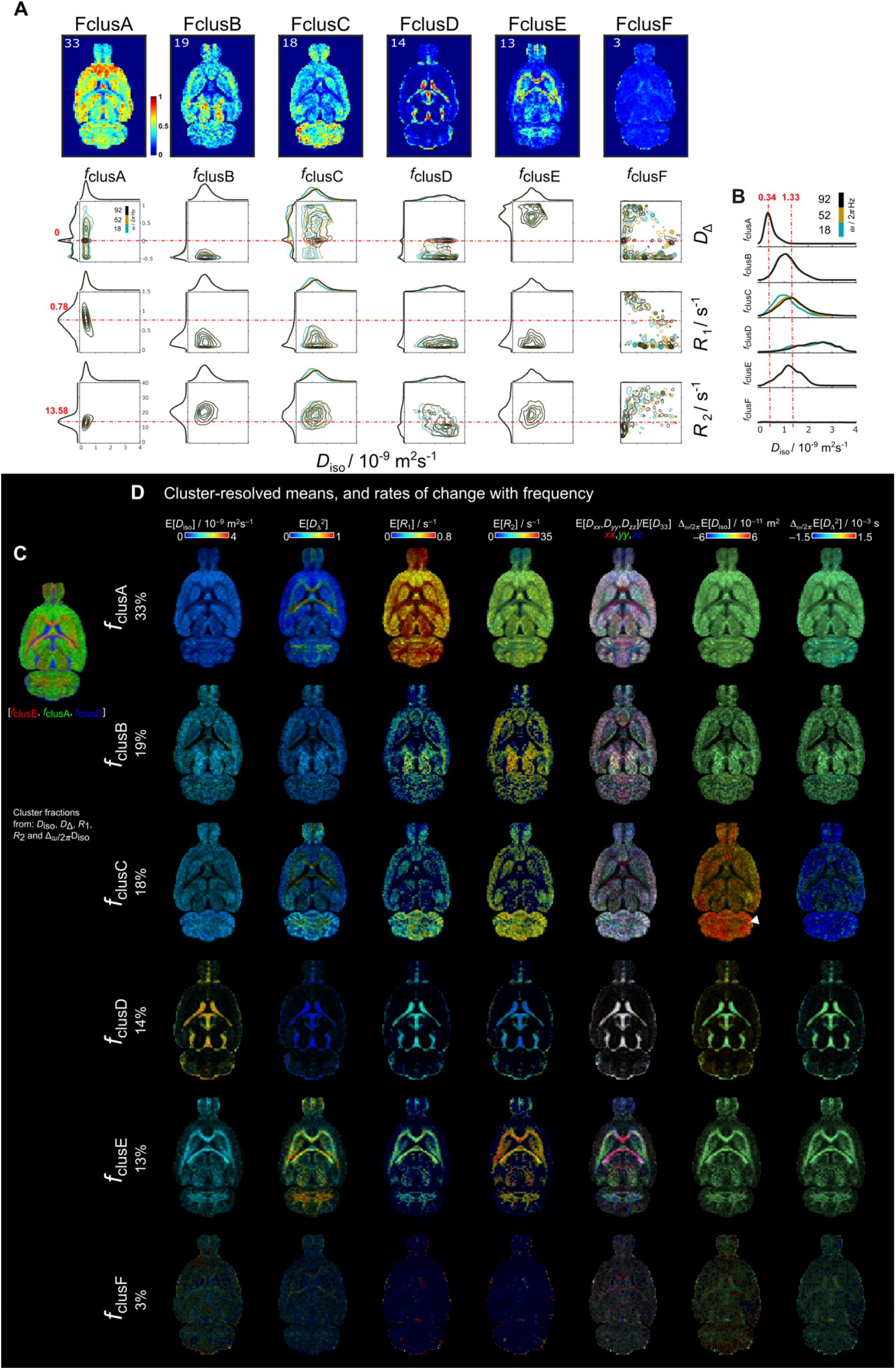
Parametric distributions and cluster-resolved mean maps of *in vivo* rat brain derived from *k* = 6 cluster-fractions including *D*_iso_, *D*_Δ_, *R*_1_, *R*_2_, and Δ_ω/2π_*D*_iso_ into the clustering. **(A)** Per-voxel cluster frequency (columns) ordered from more percentage of data included into the cluster to the least. Contour plots of the *D*_iso_-*D*_Δ_, *D*_iso_-*R*_1_, and *D*_iso_-*R*_2_ projections (rows) of the **D**(*ω*)-*R*_1_-*R*_2_ distributions for each cluster. Contour plots have different color according to ω/2π (blue = 18, mustard = 52 and black = 92 Hz). The dashed red lines indicate the mode of the *D*_Δ_, *R*_1_, and *R*_2_ parameters distributions from clusA. **(B)** Distribution amplitude of *D*_iso_ dimension displays the comparison across cluster-fractions. **(C)** Unsupervised cluster-fractions *f*_clusE_, *f*_clusA_ and *f*_clusD_ in RGB color, respectively. This representation highlights the clusters-fractions that are WM, GM and FW-like. **(D)** Cluster-resolved mean parametric maps for clusC and clusE. Arrowhead indicates the cerebellum. The cluster resolved maps were calculated using the **D**(*ω*)-*R*_1_-*R*_2_ distributions at a frequency *ω*/2π = 18 Hz.

For plotting the frequency dependance, the centroid frequencies values at 10, 50 and 90% (*ω* /2π = 18, 52 and 92 Hz) were used. Contour plots for each fraction (**Figure 7A**) showed that *f*_clusA_ was mainly characterized by the mode on *D*_Δ_ = 0. The cluster fraction *f*_clusB_, had higher *D*_iso_ and mode of *D*_Δ_ = -0.5 compared to *f*_clusB_. A clear frequency dependance in both *D*_iso_ and *D*_Δ_ was displayed in *f*_clusC_. The cluster fraction considered as the FW-like fraction, *f*_clusD_, displayed the highest values of *D*_iso_ > ∼1.5 ×10^-9^ m^2^s^-1^ together with low *R*_1_ and *R*_2_. Regarding the WM-like fraction *f*_clusE_, it contained the highest *D*_Δ_ > ∼0.5 and high *R*_2_. Cluster-resolved parameter maps (**Figure 7D**) of *f*_clusC_ and *f*_clusE_ allowed to visualize that the highest values of Δ_ω/2π_E[*D*_iso_] in *f*_clusC_ were in the cerebellum whereas *f*_clusE_ exhibited high values of E[*D*_Δ_] and E[*R*_2_] in WM regions.

In the *in vivo* human brain, the parameters included into the clustering were *D*_iso_, *D*_Δ_, *R*_1_ and *R*_2_. The parameter Δ_ω/2π_*D*_iso_ was excluded due to the limited frequency range available in the dataset (4-9 Hz). Per-voxel cluster frequency was obtained using *k* = 2 to *k* =10 (**Supplementary Figure 4**). The results from *k* = 6 were used to display the cluster fractions contour plots and cluster-resolved maps from a representative subject (**Figure 8**). Although the *in vivo* human data resulted in similar number of clusters as *in vivo* rat brain data, there is a clear absence of a frequency-dependent fraction. However, the clustering effectively delineated spherical and linear shapes as observed in the *D*_Δ_ parameter. *f*_clusA_ (22%) was primarily in GM regions such as cortex and deep GM characterized by low *D*_iso_ and the mode on spherical shapes *D*_Δ_ = 0. *f*_clusB_ (21%) in WM regions had low *D*_iso,_ *D*_Δ_ > 0.5, and higher *R*_2_ than *f*_clusB_. *f*_clusC_ (21%) was across WM and GM regions characterized by mode on *D*_Δ_ = -0.5. *f*_clusD_ was mainly located in ventricles and periphery of the brain (18%) with mode in *D*_iso_ ≅ 3×10^-9^ m^2^s^-1^. *f*_clusD_ was mainly located in ventricles and periphery of the brain (18%) with mode in *D*_iso_ > 3×10^-9^ m^2^s^-1^, while *f*_clusE_ had a larger range of *D*_iso_ between 1×10^-9^ m^2^s^-1^ and 4×10^-9^ m^2^s^-1^. Finally, *f*_clusF_ (4%) was found in the periphery of the brain.

**Figure 8.**
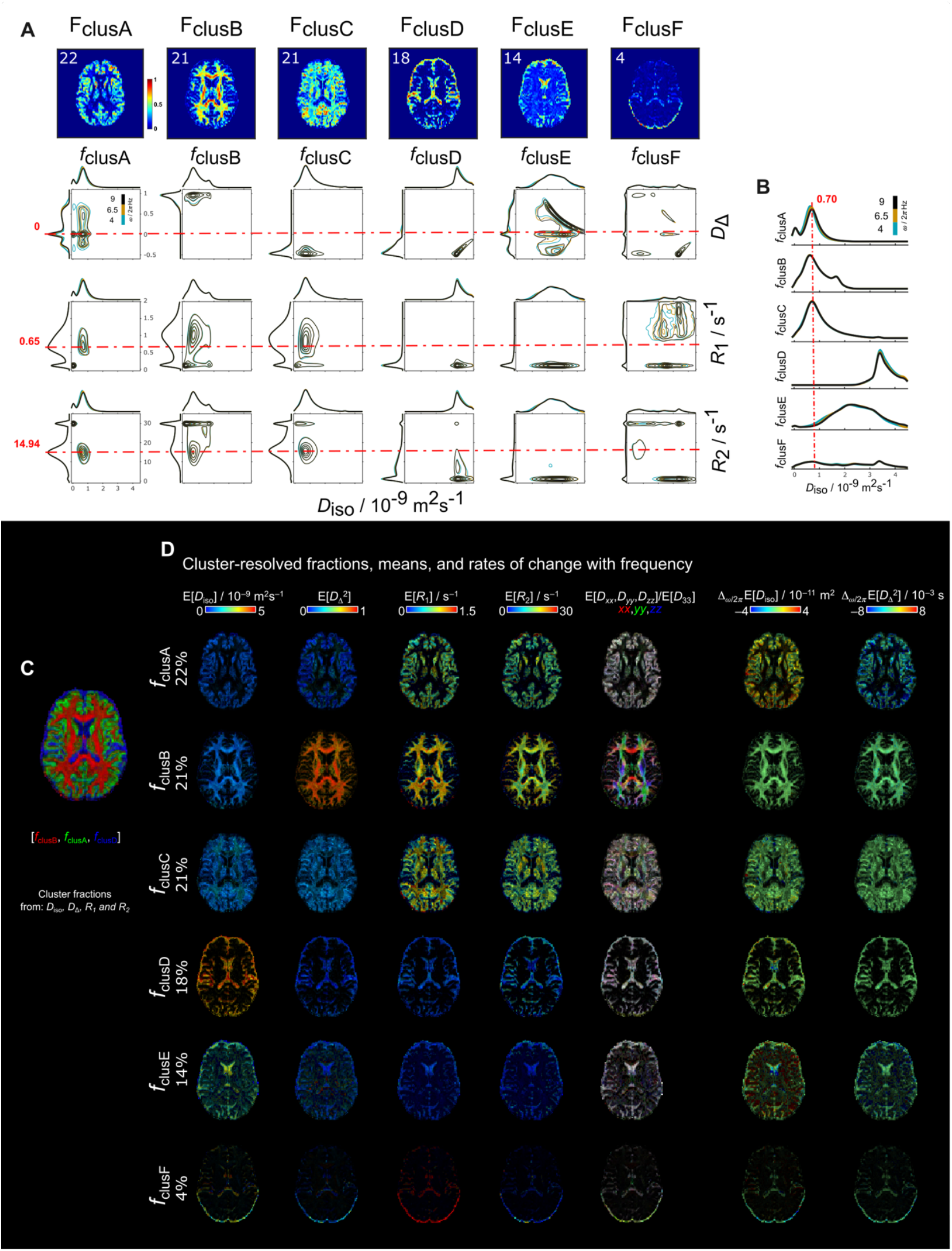
Parametric distributions and cluster-resolved mean maps of *in vivo* human brain derived from *k* = 6 cluster-fractions including *D*_iso_, *D*_Δ_, *R*_1_, and *R*_2_ into the clustering. **(A)** Per-voxel cluster frequency (columns) ordered from more percentage of data included into the cluster to the least. Contour plots of the *D*_iso_-*D*_Δ_, *D*_iso_-*R*_1_, and *D*_iso_-*R*_2_ projections (rows) of the **D**(*ω*)-*R*_1_-*R*_2_ distributions for each cluster. Contour plots have different color according to ω/2π (blue = 4, mustard = 6.5 and black = 9 Hz). **(B)** Distribution amplitude of *D*_iso_ dimension displays the comparison across cluster-fractions. **(C)** Unsupervised cluster-fractions *f*_clusB_, *f*_clusA_ and *f*_clusD_ in RGB color, respectively. **(D)** Cluster-resolved mean parametric maps. The cluster-fractions are ordered from higher to lower percentage of data belonging to each cluster. The cluster resolved maps were calculated using the **D**(*ω*)-*R*_1_-*R*_2_ distributions at a frequency *ω*/2π = 4 Hz.

We computed the clusters for all the 6 human subjects to test repeatability (**Figure 9**). For all the subjects the clusters were calculated independently and with *k* = 2-10. We found that at *k* = 6, the per-voxel cluster frequencies were similar across subjects. Hence, we plot the clusters centroids and covariance matrices in the in the normalized *D*_iso_-*D*_Δ_ space for two out of the six clusters (**Figure 9A**). The selected clusters to display were those in WM and GM regions. For the WM clusters, normalized *D*_Δ_ across subjects were approximately on 1.6 with some variation of *D*_iso_ around 0.7 and 0.9. The GM clusters displayed more consistent centroid on the *D*_iso_ dimension around 0.65. We observed that the WM and GM cluster fractions (**Figure 9B**) have a similar mode across subjects. Additionally, the *D*_iso_ and *D*_Δ_^2^ cluster-resolved maps of the WM and GM clusters are displayed from subject 1 **(Figure 9C**).

**Figure 9.**
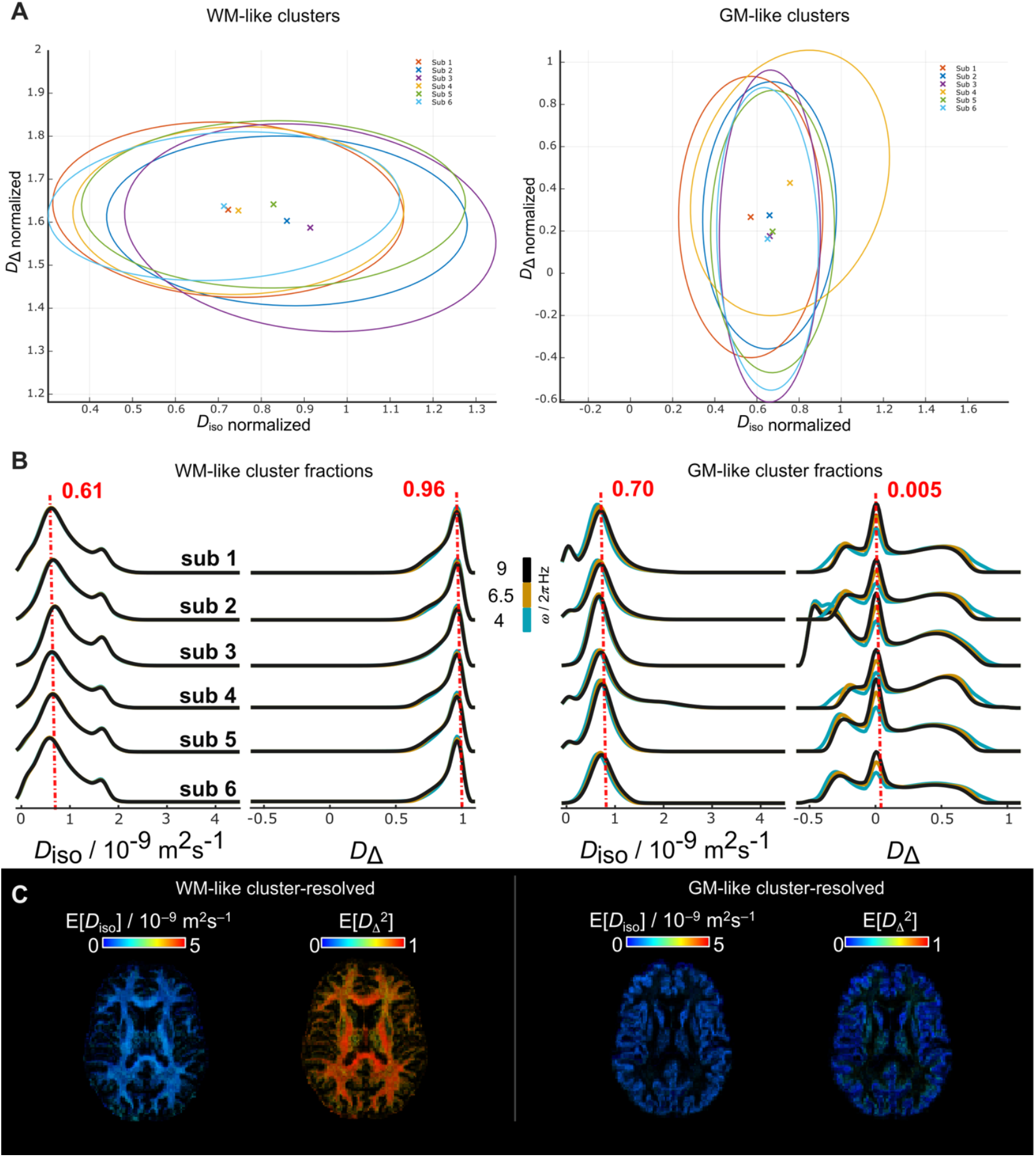
WM- and GM-like cluster-fractions of *in vivo* human brains derived from k = 6 including *D*_iso_, *D*_Δ_, *R*_1_, and *R*_2_ into the clustering. **(A)** Centroids and covariances of the WM- and GM-like clusters for each subject derived from *k = 6*. The clusters are displayed in the normalized *D*_iso_*-D*_Δ_ space.**(B)** WM- and GM-like cluster fractions of the *D*_iso_ and *D*_Δ_ parameters across all the subjects. The color of the lines indicates different ω/2π (blue = 4, mustard = 6.5 and black = 9 Hz). **(C)** Respective cluster-resolved E[*D*_iso_] and E[*D*_Δ_] maps of the WM- and GM-like fractions from sub 1.

## 4 Discussion

In this study, we identified distinct water populations from nonparametric **D**(*ω*)-*R*_1_-*R*_2_-distributions using unsupervised clustering. Our method was evaluated in both *in vivo* and *ex vivo* datasets. The number of discernible clusters was influenced by the acquisition protocol, with broader relaxation and frequency-dependent parameters enabling more detailed clustering outcomes. Despite protocol differences, at least four biological relevant fractions—WM-like, two GM-like, and FW-like—were consistently observed across healthy human and rat *in vivo* brain data, as well as *ex vivo* rat brain. This flexible, data-driven approach offers broad potential for application to other tissues, including brain tumors, heart, prostate, and breast.

Rather than depending on single per-voxel trace and anisotropy values to separate water populations, bin division of non-parametric multidimensional diffusion-relaxation MRI uses the per-voxel *D*_iso_-*D*_Δ_^2^ distribution space to differentiate WM, GM and FW contribution to a voxel. Additionally, it has been observed the advantages of adding dimensions such as in (De Almeida Martins *et al*., 2020), in which the manual binning of the *D*_iso_-*D*_Δ_^2^-*R*_2_ space allowed the distinction of cortical GM and deep GM regions. This technique facilitates the differentiation of various water populations in a single voxel, as well as the assessment of their relative contributions. Nonetheless, bin limits are still selected manually and required adjustment depending on whether the study involves *in vivo* or *ex vivo* tissue, as well as if the data type varies from brain tissue.

Clustering methods address the limitations of manual bin selection by providing an unbiased, data-driven approach. Unsupervised methods using tensor distribution data have improved the characterization of diffuse gliomas (Song *et al*., 2022). Simultaneous diffusion and relaxation measurements enhance water population differentiation, aiding astrogliosis analysis (Benjamini *et al*., 2023) and cortical layers differentiation (Kundu *et al*., 2023). In addition to relaxation and diffusion parameters, our study incorporates restriction properties characterized by the frequency-dependent domain in the non-parametric multidimensional distributions. Considering variable diffusion time regimes enables a more thorough assessment of tissue microstructure. Previous work has demonstrated that *in vivo* rat GM can contain multiple water populations, each exhibiting distinct restriction properties, reflecting the heterogeneity of the tissue environment (Yon *et al*., 2024).

The incorporation of relaxation and frequency dependence significantly enhances the ability to resolve tissue heterogeneity within a voxel. In this context, applying the unsupervised approach to the non-parametric **D**(*ω*)-*R*_1_-*R*_2_-distributions enables the extraction of per-voxel distributions of multiple water populations with greater biological relevance, as evidenced by their correspondence to histological counterparts. In contrast, relying on manual *D*_iso_ and *D*_Δ_^2^ bin-limits in high-dimensional data can lead to the mixing of tissue fractions that are unlikely to biologically coexist within the same spatial domain. For example, in the manual bin approach, WM-like fraction (bin1) exhibited high Δ_*ω*/2π_*D*_iso_ values within the granule cell layer. This ambiguity was resolved through unsupervised clustering, which correctly assigned the granule cell layer to a separate tissue fraction. A similar challenge arises in GM, where variations in cellular density, cell size, axonal content, and chemical composition differ across distinct brain regions. The unsupervised clustering approach provided characteristic cluster fraction distributions, offering a more accurate and biologically meaningful delineation of tissue heterogeneity.

We applied the unsupervised clustering approach to both *in vivo* and *ex vivo* datasets, revealing differences in the number of clusters identified between *ex vivo* and *in vivo* rat and *in vivo* human brains. These differences were primarily attributed to variations in acquisition protocols. The *ex vivo* settings allowed for higher spatial resolution and the collection of more extensive data due to the longer acquisition times, whereas *in vivo* acquisitions were constrained by shorter imaging sessions, MRI gradient strength, and motion-related limitations. Despite these protocol-dependent differences, we consistently identified similar biologically relevant fractions across both *ex vivo* and *in vivo* datasets. A recent work demonstrated the reliability and reproducibility of non-parametric **D**(*ω*)-*R*_1_-*R*_2_-distributions in the human brain (Manninen *et al*., 2024). Additionally, our repeatability experiments showed that unsupervised clustering within the same protocol and acquisition settings results in similar centroids and covariance matrices.

Notably, certain components were relatively straightforward to interpret, such as the high *D*_Δ_ distributions, which were predominantly associated with WM regions. In contrast, the zero and negative anisotropic distributions without frequency-dependence, were more challenging to interpret biologically. However, these components were notably more prominent in GM, suggesting underlying variations in cellular and microstructural properties. When frequency dependence data were available, as in the *ex vivo* and *in vivo* rat datasets, the clustering approach enabled the extraction of a fraction resembling the apparent intra-soma compartment. This fraction was characterized by increased *D*_iso_ at higher frequencies, consistent with previous findings using oscillating gradient spin echo acquisitions (OGSE) (Does *et al*., 2003; Aggarwal *et al*., 2012; Baron *et al*., 2014; Oshiro *et al*., 2024). Due to the limited frequency range in the *in vivo* human data, this intra-soma-like feature could not be resolved, this could be addressed by using clinical MRI scanner with stronger gradients (Tan *et al*., 2020; Hennel *et al*., 2021; Xu, 2021; Dai *et al*., 2023).

The clusters obtained in this study should not be interpreted as isolated features corresponding to a single microstructural property. For instance, a cluster characterized by high Δ_ω/2π_*D*_iso_ may be attributed to the apparent soma fraction, analogously to the time-dependent microscopic kurtosis (μK) (Novello *et al*., 2022). However, it is essential to acknowledge that the pronounced restriction properties revealed by the frequency dependence of diffusion can also reflect underlying exchange effects. This phenomenon has been demonstrated by Chakwizira *et al*., who showed that the sensitivity to restriction or exchange can be modulated by varying the design of gradient waveforms (Chakwizira *et al*., 2023). In the current study, we employed a broad spectrum of gradient waveforms, which inherently introduces the potential for confounding effects between restriction and exchange. To disentangle these contributions more effectively, a targeted approach could be considered, such as selecting a subset of the data with waveforms specifically optimized to enhance sensitivity to either restriction or exchange.

The inclusion of frequency dependence is crucial for probing the microstructure of GM microstructure. Specifically, for Δ_ω/2π_*D*_Δ_, we observed that increasing the frequency shifts the anisotropy distribution closer to 0. A tendency for FA to slightly decrease at lower diffusion times has been previously noted in time and frequency-dependent diffusion studies (Tan *et al*., 2020; Tétreault *et al*., 2020; Ba *et al*., 2024). However, in contrast with FA and μFA, *D*_Δ_ distributions provides a more detailed representation of tensor shapes, distinguishing between prolate and oblate (Lawrenz *et al*., 2010; Shemesh *et al*., 2016b). In our observations, at low frequencies, the Δ_ω/2π_*D*_Δ_ values in GM regions are predominantly closer to 1 (stick-like structures) and -0.5. As the frequency increases, a distinct shift occurs in the distribution toward *D*_Δ_ = 0 (spherical structures). This observation highlights the value of frequency dependence in identifying and characterizing specific microstructural compartments within GM. Investigating the soma more precisely—whether through biophysical modeling or signal representation—requires selecting an optimal frequency or time-dependent range. Intermediate ranges may incorporate contributions from other microstructural features, thereby complicating the interpretation and potentially leading to suboptimal signal fitting. There are only few studies about the time-dependent effect in the microscopic anisotropy (Ianuş *et al*., 2018; Narvaez *et al*., 2022, 2024; Johnson *et al*., 2024; Yon *et al*., 2025).

The use of Monte Carlo inversion algorithms to resolve multidimensional parameter spaces is a well-established method (Prange *et al*., 2009), supported by prior studies and simulations (De Almeida Martins *et al*., 2018; Narvaez *et al*., 2024). This approach explores the entire solution space through multiple component decomposition, which, while comprehensive, can introduce challenges such as data overfitting and the generation of highly complex distributions. To address potential overfitting, the resulting solutions are typically weighted and summarized into statistical descriptors, such as medians, variances, and covariances, to enable the calculation of parametric maps (Yon *et al*., 2020; De Almeida Martins *et al*., 2021; Martin *et al*., 2021; Narvaez *et al*., 2022, 2024). In the current study, we leveraged the weighted solutions directly as input for clustering, acknowledging that not all data points contribute equally to the overall biological relevance of the solution space. By doing so, we aimed to retain the granularity of the multidimensional data while reducing the influence of less significant components. The Gaussian Mixture Model (GMM) first iteratively determines the centroids (μ) of the clusters, with each centroid representing a meaningful data point. In contrast, the covariance (σ) reflects the spread and shape of the clusters, which plays a key role in preventing data overfitting. When applied to non-parametric **D**(*ω*)-*R*_1_-*R*_2_-distributions, these GMM parameters correspond to the mean and variance of the distributions, both of which require careful interpretation. As the number of clusters increases within a given tissue, additional cluster fractions with less than approximately 8% of the data start to emerge. To avoid overfitting, we limited the number of clusters selected for each experiment. The validity of the identified cluster fractions can only be confirmed through spatial correlation with well-established biological structures, which necessitates histological comparisons for accurate biological interpretation.

In this study, we utilized two histological stainings, Nissl staining for cell bodies and myelin staining to identify myelinated fibers. While these stainings provide valuable insight into cellular density and myelin distribution, they do not capture the full complexity of the tissue microstructure. Brain tissue consists of diverse components, including various cell populations, dendrites, unmyelinated axons, vasculature, and extracellular matrix, which were not accounted for in this study, thus limiting the interpretation of the cluster fractions. Furthermore, while light microscopy is a well-established tool, its resolution is limited to visualize certain microstructural features, such as subcellular components and neuropil. These features are critical for understanding the full extent of tissue heterogeneity, particularly in GM regions with diverse cellular, axonal and synaptic characteristics. To strengthen the associations between MRI and histology, more comprehensive analyses incorporating additional stainings (*e*.*g*., for neuronal and glial markers, vasculature, or extracellular components) and advanced imaging techniques are needed. Electron microscopy (EM) could play a crucial role in this process, providing much higher resolution images that allow for visualization of ultrastructural features, such as synaptic junctions, fine axonal structures, and the organization of myelin at a sub-micrometer scale. Such insights would significantly enhance our understanding of the tissue microstructure and improve the biological interpretation of MRI data. Moreover, the integration of advanced image analysis techniques, including 3D reconstructions, machine learning-based segmentation, and quantitative morphometric analyses, could offer more detailed and precise evaluations of tissue organization (Abdollahzadeh *et al*., 2021; Larsen *et al*., 2021; Salo *et al*., 2021). Future work should thus incorporate these complementary methods to provide a more comprehensive view of tissue organization and its relationship to multidimensional MRI-based measures.

## 5 Conclusions and Outlook

In conclusion, this study highlights the potential of unsupervised classification of multidimensional diffusion-relaxation correlation data to enhance tissue characterization at the voxel level. The method effectively disentangles tissue components and provides valuable insights into diffusivity, anisotropy, and chemical properties, which are crucial for identifying tissue changes in pathologies such as tumors, demyelination, Wallerian degeneration, and cell swelling. This approach offers significant advantages over traditional methods, enabling a more comprehensive understanding of tissue microstructure without relying on assumptions. While promising, further validation is needed in clinical settings, particularly with complementary histological data. Expanding this method to other tissues and diseases could broaden its applications, making it a powerful tool for both research and clinical diagnostics, especially in neuroimaging and other medical fields.

## Author Contributions

ON: conceptualization, methodology, data acquisition and processing, data analysis, visualization, manuscript writing, review, and editing. MY: methodology, data acquisition and processing, manuscript review and editing. RS: conceptualization, data analysis. JK: methodology, data acquisition and processing. ME: data processing and analysis. EP: acquiring license and imaging permits, data acquisition and processing. VL: acquiring license and imaging permits. JH: acquiring license and imaging permits, and radiological reviewing. FL: data acquisition. DT: methodology, manuscript review, and editing. AS: conceptualization, funding acquisition, project administration, supervision, manuscript writing, review, and revision. All authors revised and contributed to the final version of the manuscript.

## Acknowledgments

This work was carried out with the support of Kuopio Biomedical Imaging Unit, University of Eastern Finland, Kuopio, Finland (part of Finnish Biomedical Imaging Node, EuroBioImaging). The computational analyzes were performed on servers provided by UEF Bioinformatics Center, University of Eastern Finland, Finland and CSC – IT Center for Science, Finland. The authors would like to thank Maarit Pulkkinen for her help with the animal handling.

## Data and Code Availability

MATLAB source code for preprocessing, Monte-Carlo data inversion and clustering is freely available at https://github.com/UEF-Multiscale-Imaging. The data used in this study is available upon request.

## References

Abdollahzadeh, A., Belevich, I., Jokitalo, E., Sierra, A. and Tohka, J. (2021) “DeepACSON automated segmentation of white matter in 3D electron microscopy,” Communications Biology, 4(1), pp. 1–14. Available at: 10.1038/s42003-021-01699-w.

Aggarwal, M. (2020) “Chapter 4. Restricted Diffusion and Spectral Content of the Gradient Waveforms,” in D. Topgaard (ed.) New Developments in NMR. Cambridge: Royal Society of Chemistry, pp. 103–122. Available at: 10.1039/9781788019910-00103.

Aggarwal, M., Jones, M.V., Calabresi, P.A., Mori, S. and Zhang, J. (2012) “Probing mouse brain microstructure using oscillating gradient diffusion MRI,” Magnetic Resonance in Medicine, 67(1), pp. 98–109. Available at: 10.1002/mrm.22981.

Aggarwal, M., Smith, M.D. and Calabresi, P.A. (2020) “Diffusion-time dependence of diffusional kurtosis in the mouse brain,” Magnetic Resonance in Medicine, 84(3), pp. 1564– 1578. Available at: 10.1002/mrm.28189.

Andersson, J.L.R., Skare, S. and Ashburner, J. (2003) “How to correct susceptibility distortions in spin-echo echo-planar images: application to diffusion tensor imaging,” NeuroImage, 20(2), pp. 870–888. Available at: 10.1016/S1053-8119(03)00336-7.

Arbabi, A., Kai, J., Khan, A.R. and Baron, C.A. (2020) “Diffusion dispersion imaging: Mapping oscillating gradient spin-echo frequency dependence in the human brain,” Magnetic Resonance in Medicine, 83(6), pp. 2197–2208. Available at: 10.1002/mrm.28083.

Ba, R., Kang, L. and Wu, D. (2024) “Time-dependent diffusion magnetic resonance imaging: measurement, modeling, and applications,” Journal of Zhejiang University-SCIENCE A, 25(10), pp. 765–787. Available at: 10.1631/jzus.A2400139.

Bankhead, P., Loughrey, M.B., Fernández, J.A., Dombrowski, Y., McArt, D.G., Dunne, P.D., et al. (2017) “QuPath: Open source software for digital pathology image analysis,” Scientific Reports, 7(1), p. 16878. Available at: 10.1038/s41598-017-17204-5.

Baron, C.A. and Beaulieu, C. (2014) “Oscillating gradient spin-echo (OGSE) diffusion tensor imaging of the human brain,” Magnetic Resonance in Medicine, 72(3), pp. 726–736. Available at: 10.1002/mrm.24987.

Basser, P.J., Mattiello, J. and LeBihan, D. (1994) “MR diffusion tensor spectroscopy and imaging,” Biophysical Journal, 66(1), pp. 259–267. Available at: 10.1016/S0006-3495(94)80775-1.

Benjamini, D. and Basser, P.J. (2017) “Magnetic resonance microdynamic imaging reveals distinct tissue microenvironments,” NeuroImage, 163, pp. 183–196. Available at: 10.1016/j.neuroimage.2017.09.033.

Benjamini, D., Priemer, D.S., Perl, D.P., Brody, D.L. and Basser, P.J. (2023) “Mapping astrogliosis in the individual human brain using multidimensional MRI,” Brain, 146(3), pp. 1212–1226. Available at: 10.1093/brain/awac298.

Budde, M.D. and Frank, J.A. (2012) “Examining brain microstructure using structure tensor analysis of histological sections,” NeuroImage, 63(1), pp. 1–10. Available at: 10.1016/j.neuroimage.2012.06.042.

Chakwizira, A., Zhu, A., Foo, T., Westin, C.-F., Szczepankiewicz, F. and Nilsson, M. (2023) “Diffusion MRI with free gradient waveforms on a high-performance gradient system: Probing restriction and exchange in the human brain,” NeuroImage, 283, p. 120409. Available at: 10.1016/j.neuroimage.2023.120409.

Chen, J., Ades-Aron, B., Lee, H.-H., Mehrin, S., Pang, M., Novikov, D.S., et al. (2024) “Optimization and validation of the DESIGNER preprocessing pipeline for clinical diffusion MRI in white matter aging,” Imaging Neuroscience, 2, pp. 1–17. Available at: 10.1162/imag_a_00125.

Coelho, S., Baete, S.H., Lemberskiy, G., Ades-Aron, B., Barrol, G., Veraart, J., et al. (2022) “Reproducibility of the Standard Model of diffusion in white matter on clinical MRI systems,” NeuroImage, 257, p. 119290. Available at: 10.1016/j.neuroimage.2022.119290.

Dai, E., Zhu, A., Yang, G.K., Quah, K., Tan, E.T., Fiveland, E., et al. (2023) “Frequency-dependent diffusion kurtosis imaging in the human brain using an oscillating gradient spin echo sequence and a high-performance head-only gradient,” NeuroImage, 279, p. 120328. Available at: 10.1016/j.neuroimage.2023.120328.

De Almeida Martins, J.P., Tax, C.M.W., Reymbaut, A., Szczepankiewicz, F., Chamberland, M., Jones, D.K., et al. (2021) “Computing and visualising intra-voxel orientation-specific relaxation–diffusion features in the human brain,” Human Brain Mapping, 42(2), pp. 310–328. Available at: 10.1002/hbm.25224.

De Almeida Martins, J.P., Tax, C.M.W., Szczepankiewicz, F., Jones, D.K., Westin, C.-F. and Topgaard, D. (2020) “Transferring principles of solid-state and Laplace NMR to the field of in vivo brain MRI,” Magnetic Resonance, 1(1), pp. 27–43. Available at: 10.5194/mr-1-27-2020.

De Almeida Martins, J.P. and Topgaard, D. (2018) “Multidimensional correlation of nuclear relaxation rates and diffusion tensors for model-free investigations of heterogeneous anisotropic porous materials,” Scientific Reports, 8(1), p. 2488. Available at: 10.1038/s41598-018-19826-9.

Does, M.D., Parsons, E.C. and Gore, J.C. (2003) “Oscillating gradient measurements of water diffusion in normal and globally ischemic rat brain,” Magnetic Resonance in Medicine, 49(2), pp. 206–215. Available at: 10.1002/mrm.10385.

Hennel, F., Michael, E.S. and Pruessmann, K.P. (2021) “Improved gradient waveforms for oscillating gradient spin-echo (OGSE) diffusion tensor imaging,” NMR in Biomedicine, 34(2), p. e4434. Available at: 10.1002/nbm.4434.

Henriques, R.N., Palombo, M., Jespersen, S.N., Shemesh, N., Lundell, H. and Ianuş, A. (2021) “Double diffusion encoding and applications for biomedical imaging,” Journal of Neuroscience Methods, 348, p. 108989. Available at: 10.1016/j.jneumeth.2020.108989.

Ianus, A., Jespersen, S.N., Serradas Duarte, T., Alexander, D.C., Drobnjak, I. and Shemesh, N. (2018) “Accurate estimation of microscopic diffusion anisotropy and its time dependence in the mouse brain,” NeuroImage, 183, pp. 934–949. Available at: 10.1016/j.neuroimage.2018.08.034.

Jelescu, I.O., Palombo, M., Bagnato, F. and Schilling, K.G. (2020) “Challenges for biophysical modeling of microstructure,” Journal of Neuroscience Methods, 344, p. 108861. Available at: 10.1016/j.jneumeth.2020.108861.

Jiang, H., Svenningsson, L. and Topgaard, D. (2023) “Multidimensional encoding of restricted and anisotropic diffusion by double rotation of the q vector,” Magnetic Resonance, 4(1), pp. 73–85. Available at: 10.5194/mr-4-73-2023.

Johnson, J.T.E., Irfanoglu, M.O., Manninen, E., Ross, T.J., Yang, Y., Laun, F.B., et al. (2024) “In vivo disentanglement of diffusion frequency-dependence, tensor shape, and relaxation using multidimensional MRI,” Human Brain Mapping, 45(7), p. e26697. Available at: 10.1002/hbm.26697.

Kellner, E., Dhital, B., Kiselev, V.G. and Reisert, M. (2016) “Gibbs-ringing artifact removal based on local subvoxel-shifts,” Magnetic Resonance in Medicine, 76(5), pp. 1574–1581. Available at: 10.1002/mrm.26054.

Kim, D., Doyle, E.K., Wisnowski, J.L., Kim, J.H. and Haldar, J.P. (2017) “Diffusion-Relaxation Correlation Spectroscopic Imaging (DR-CSI): A Multidimensional Approach for Probing Microstructure,” Magnetic resonance in medicine, 78(6), pp. 2236–2249. Available at: 10.1002/mrm.26629.

Kundu, S., Barsoum, S., Ariza, J., Nolan, A.L., Latimer, C.S., Keene, C.D., et al. (2023) “Mapping the individual human cortex using multidimensional MRI and unsupervised learning,” Brain Communications, 5(6), p. fcad258. Available at: 10.1093/braincomms/fcad258.

Laitinen, T., Sierra, A., Pitkänen, A. and Gröhn, O. (2010) “Diffusion tensor MRI of axonal plasticity in the rat hippocampus,” NeuroImage, 51(2), pp. 521–530. Available at: 10.1016/j.neuroimage.2010.02.077.

Larsen, N.Y., Li, X., Tan, X., Ji, G., Lin, J., Rajkowska, G., et al. (2021) “Cellular 3D-reconstruction and analysis in the human cerebral cortex using automatic serial sections,” Communications Biology, 4(1), pp. 1–15. Available at: 10.1038/s42003-021-02548-6.

Lasic, S., Szczepankiewicz, F., Eriksson, S., Nilsson, M. and Topgaard, D. (2014) “Microanisotropy imaging: quantification of microscopic diffusion anisotropy and orientational order parameter by diffusion MRI with magic-angle spinning of the q-vector,” Frontiers in Physics, 2. Available at: 10.3389/fphy.2014.00011.

Lawrenz, M., Koch, M.A. and Finsterbusch, J. (2010) “A tensor model and measures of microscopic anisotropy for double-wave-vector diffusion-weighting experiments with long mixing times,” Journal of Magnetic Resonance, 202(1), pp. 43–56. Available at: 10.1016/j.jmr.2009.09.015.

Lundell, H., Nilsson, M., Dyrby, T.B., Parker, G.J.M., Cristinacce, P.L.H., Zhou, F.-L., et al. (2019) “Multidimensional diffusion MRI with spectrally modulated gradients reveals unprecedented microstructural detail,” Scientific Reports, 9(1), p. 9026. Available at: 10.1038/s41598-019-45235-7.

Manninen, E., Bao, S., Landman, B.A., Yang, Y., Topgaard, D. and Benjamini, D. (2024) “Variability of multidimensional diffusion–relaxation MRI estimates in the human brain,” Imaging Neuroscience, 2, pp. 1–24. Available at: 10.1162/imag_a_00387.

Martin, J., Endt, S., Wetscherek, A., Kuder, T.A., Doerfler, A., Uder, M., et al. (2020) “Contrast-to-noise ratio analysis of microscopic diffusion anisotropy indices in q-space trajectory imaging,” Zeitschrift für Medizinische Physik, 30(1), pp. 4–16. Available at: 10.1016/j.zemedi.2019.01.003.

Martin, J., Reymbaut, A., Schmidt, M., Doerfler, A., Uder, M., Laun, F.B., et al. (2021) “Nonparametric D-R1-R2 distribution MRI of the living human brain,” NeuroImage, 245, p. 118753. Available at: 10.1016/j.neuroimage.2021.118753.

Molina, I.S.M., Salo, R.A., Abdollahzadeh, A., Tohka, J., Gröhn, O. and Sierra, A. (2020) “In Vivo Diffusion Tensor Imaging in Acute and Subacute Phases of Mild Traumatic Brain Injury in Rats,” eNeuro, 7(3). Available at: 10.1523/ENEURO.0476-19.2020.

Narvaez, O., Svenningsson, L., Yon, M., Sierra, A. and Topgaard, D. (2022) “Massively Multidimensional Diffusion-Relaxation Correlation MRI,” Frontiers in Physics, 9, p. 793966. Available at: 10.3389/fphy.2021.793966.

Narvaez, O., Yon, M., Jiang, H., Bernin, D., Forssell-Aronsson, E., Sierra, A., et al. (2024) “Nonparametric distributions of tensor-valued Lorentzian diffusion spectra for model-free data inversion in multidimensional diffusion MRI,” The Journal of Chemical Physics, 161(8), p. 084201. Available at: 10.1063/5.0213252.

Nilsson, M., Szczepankiewicz, F., Lampinen, B., Ahlgren, A., De Almeida Martins, J.P., Lasic, S., et al. (2018) “An open-source framework for analysis of multidimensional diffusion MRI data implemented in MATLAB,” in International Society of Magnetic Resonance in Medicine. Paris, France: International Society of Magnetic Resonance in Medicine.

Novello, L., Henriques, R.N., Ianuş, A., Feiweier, T., Shemesh, N. and Jovicich, J. (2022) “In vivo Correlation Tensor MRI reveals microscopic kurtosis in the human brain on a clinical 3T scanner,” NeuroImage, 254, p. 119137. Available at: 10.1016/j.neuroimage.2022.119137.

Novikov, D.S., Kiselev, V.G. and Jespersen, S.N. (2018) “On modeling,” Magnetic Resonance in Medicine, 79(6), pp. 3172–3193. Available at: 10.1002/mrm.27101.

Novikov, D.S., Veraart, J., Jelescu, I.O. and Fieremans, E. (2018) “Rotationally-invariant mapping of scalar and orientational metrics of neuronal microstructure with diffusion MRI,” NeuroImage, 174, pp. 518–538. Available at: 10.1016/j.neuroimage.2018.03.006.

Oshiro, H., Hata, J., Nakashima, D., Oshiro, R., Hayashi, N., Haga, Y., et al. (2024) “Restricted diffusion characteristics in oscillating gradient spin echo with mesoscopic phantom,” Heliyon, 10(4). Available at: 10.1016/j.heliyon.2024.e26391.

Palombo, M., Ianus, A., Guerreri, M., Nunes, D., Alexander, D.C., Shemesh, N., et al. (2020) “SANDI: A compartment-based model for non-invasive apparent soma and neurite imaging by diffusion MRI,” NeuroImage, 215, p. 116835. Available at: 10.1016/j.neuroimage.2020.116835.

Pas, K., Komlosh, M.E., Perl, D.P., Basser, P.J. and Benjamini, D. (2020) “Retaining information from multidimensional correlation MRI using a spectral regions of interest generator,” Scientific Reports, 10(1), p. 3246. Available at: 10.1038/s41598-020-60092-5.

Pierpaoli, C., Jezzard, P., Basser, P.J., Barnett, A. and Di Chiro, G. (1996) “Diffusion tensor MR imaging of the human brain.,” Radiology, 201(3), pp. 637–648. Available at: 10.1148/radiology.201.3.8939209.

Prange, M. and Song, Y.-Q. (2009) “Quantifying uncertainty in NMR spectra using Monte Carlo inversion,” Journal of Magnetic Resonance, 196(1), pp. 54–60. Available at: 10.1016/j.jmr.2008.10.008.

Püspöki, Z., Storath, M., Sage, D. and Unser, M. (2016) “Transforms and Operators for Directional Bioimage Analysis: A Survey,” in W.H. De Vos, S. Munck, and J.-P. Timmermans (eds.) Focus on Bio-Image Informatics. Cham: Springer International Publishing (Advances in Anatomy, Embryology and Cell Biology), pp. 69–93. Available at: 10.1007/978-3-319-28549-8_3.

Reymbaut, A. (2020) “Chapter 3. Diffusion Anisotropy and Tensor-valued Encoding,” in D. Topgaard (ed.) New Developments in NMR. Cambridge: Royal Society of Chemistry, pp. 68– 102. Available at: 10.1039/9781788019910-00068.

Reymbaut, A., Critchley, J., Durighel, G., Sprenger, T., Sughrue, M., Bryskhe, K., et al. (2021) “Toward nonparametric diffusion-characterization of crossing fibers in the human brain,” Magnetic Resonance in Medicine, 85(5), pp. 2815–2827. Available at: 10.1002/mrm.28604.

Salo, R.A., Belevich, I., Jokitalo, E., Gröhn, O. and Sierra, A. (2021) “Assessment of the structural complexity of diffusion MRI voxels using 3D electron microscopy in the rat brain,” NeuroImage, 225, p. 117529. Available at: 10.1016/j.neuroimage.2020.117529.

Shemesh, N., Jespersen, S.N., Alexander, D.C., Cohen, Y., Drobnjak, I., Dyrby, T.B., et al. (2016a) “Conventions and nomenclature for double diffusion encoding NMR and MRI,” Magnetic Resonance in Medicine, 75(1), pp. 82–87. Available at: 10.1002/mrm.25901.

Shemesh, N., Jespersen, S.N., Alexander, D.C., Cohen, Y., Drobnjak, I., Dyrby, T.B., et al. (2016b) “Conventions and nomenclature for double diffusion encoding NMR and MRI,” Magnetic Resonance in Medicine, 75(1), pp. 82–87. Available at: 10.1002/mrm.25901.

Sjölund, J., Szczepankiewicz, F., Nilsson, M., Topgaard, D., Westin, C.-F. and Knutsson, H. (2015) “Constrained optimization of gradient waveforms for generalized diffusion encoding,” Journal of magnetic resonance (San Diego, Calif. : 1997), 261, pp. 157–168. Available at: 10.1016/j.jmr.2015.10.012.

Slator, P.J., Hutter, J., Palombo, M., Jackson, L.H., Ho, A., Panagiotaki, E., et al. (2019) “Combined diffusion-relaxometry MRI to identify dysfunction in the human placenta,” Magnetic Resonance in Medicine, 82(1), pp. 95–106. Available at: 10.1002/mrm.27733.

Song, Y., Ly, I., Fan, Q., Nummenmaa, A., Martinez-Lage, M., Curry, W.T., et al. (2022) “Measurement of Full Diffusion Tensor Distribution Using High-Gradient Diffusion MRI and Applications in Diffuse Gliomas,” Frontiers in Physics, 10, p. 813475. Available at: 10.3389/fphy.2022.813475.

Tan, E.T., Shih, R.Y., Mitra, J., Sprenger, T., Hua, Y., Bhushan, C., et al. (2020) “Oscillating Diffusion-Encoding with a High Gradient-Amplitude and High Slew-Rate Head-Only Gradient for Human Brain Imaging,” Magnetic resonance in medicine, 84(2), pp. 950–965. Available at: 10.1002/mrm.28180.

Tétreault, P., Harkins, K.D., Baron, C.A., Stobbe, R., Does, M.D. and Beaulieu, C. (2020) “Diffusion time dependency along the human corpus callosum and exploration of age and sex differences as assessed by oscillating gradient spin-echo diffusion tensor imaging,” NeuroImage, 210, p. 116533. Available at: 10.1016/j.neuroimage.2020.116533.

Topgaard, D. (2016) “Chapter 7. NMR Methods for Studying Microscopic Diffusion Anisotropy,” in R. Valiullin (ed.) New Developments in NMR. Cambridge: Royal Society of Chemistry, pp. 226–259. Available at: 10.1039/9781782623779-00226.

Topgaard, D. (2019) “Diffusion tensor distribution imaging,” NMR in Biomedicine, 32(5), p. e4066. Available at: 10.1002/nbm.4066.

Tournier, J.-D., Smith, R., Raffelt, D., Tabbara, R., Dhollander, T., Pietsch, M., et al. (2019) “MRtrix3: A fast, flexible and open software framework for medical image processing and visualisation,” NeuroImage, 202, p. 116137. Available at: 10.1016/j.neuroimage.2019.116137.

Xu, J. (2021) “Probing neural tissues at small scales: Recent progress of oscillating gradient spin echo (OGSE) neuroimaging in humans,” Journal of neuroscience methods, 349, p. 109024. Available at: 10.1016/j.jneumeth.2020.109024.

Xu, J., Li, H., Li, K., Harkins, K.D., Jiang, X., Xie, J., et al. (2016) “Fast and simplified mapping of mean axon diameter using temporal diffusion spectroscopy,” NMR in Biomedicine, 29(4), pp. 400–410. Available at: 10.1002/nbm.3484.

Yon, M., De Almeida Martins, J.P., Bao, Q., Budde, M.D., Frydman, L. and Topgaard, D. (2020) “Diffusion tensor distribution imaging of an in vivo mouse brain at ultrahigh magnetic field by spatiotemporal encoding,” NMR in Biomedicine, 33(11), p. e4355. Available at: 10.1002/nbm.4355.

Yon, M., Narvaez, O., Topgaard, D. and Sierra, A. (2024) “In vivo rat-brain mapping of multiple gray matter water populations using nonparametric D(ω)-R 1-R 2 distributions MRI.” Biophysics. Available at: 10.1101/2024.06.07.597866.

Yon, M., Narvaez, O., Topgaard, D. and Sierra, A. (2025) “In vivo rat brain mapping of multiple gray matter water populations using nonparametric D(ω)-R1-R2 distributions MRI,” NMR in Biomedicine, 38(1), p. e5286. Available at: 10.1002/nbm.5286.

Zhang, H., Schneider, T., Wheeler-Kingshott, C.A. and Alexander, D.C. (2012) “NODDI: Practical in vivo neurite orientation dispersion and density imaging of the human brain,” NeuroImage, 61(4), pp. 1000–1016. Available at: 10.1016/j.neuroimage.2012.03.072.

