## Supplementary information for "Data-driven classification of tissue water populations by massively multidimensional diffusion-relaxation correlation MRI"

[1] A.I. Virtanen Institute for Molecular Sciences, University of Eastern Finland, Kuopio, Finland, [2] Department of Chemistry, Lund University, Lund, Sweden, [3] Neurocenter, Kuopio University Hospital, Kuopio, Finland, [4] Institute of Clinical Medicine, University of Eastern Finland, Kuopio, Finland, [5] Neurocenter Neurosurgery, Kuopio University Hospital, Kuopio, Finland, [6] Department of Clinical Radiology, Kuopio University Hospital, Kuopio, Finland, [7] Institute of Radiology, University Hospital Erlangen, Erlangen, Germany

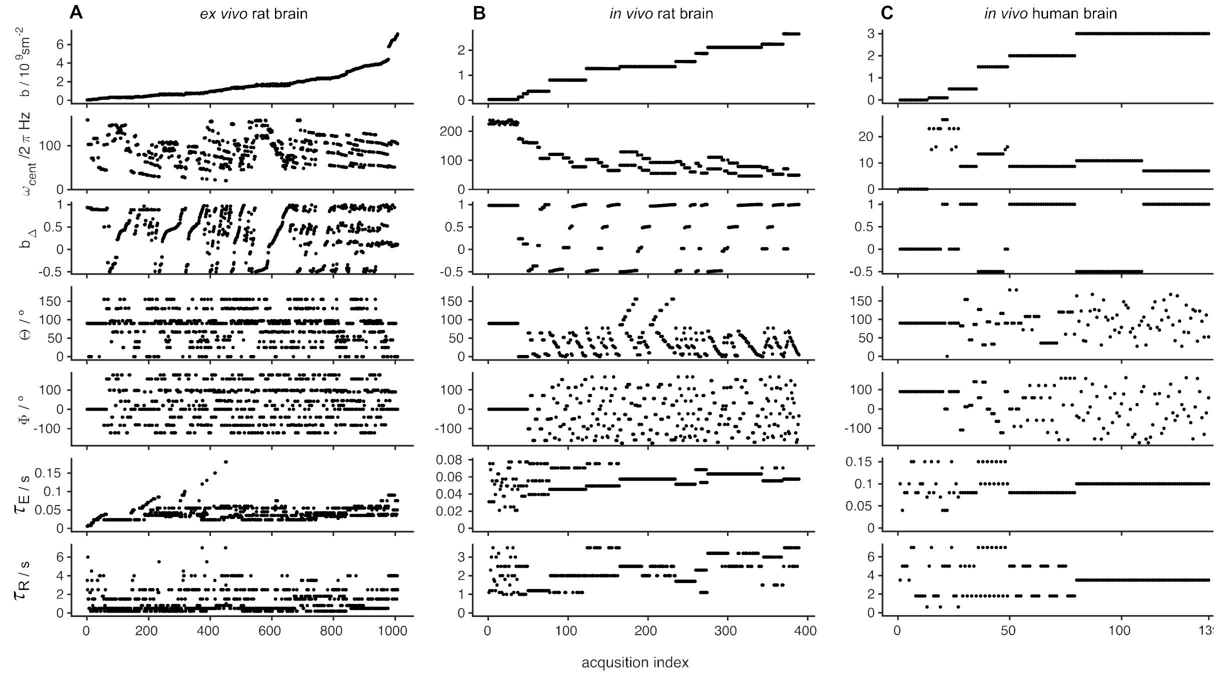


**Supplementary Figure 1.** Acquisition scheme for ex vivo rat brain (A), in vivo rat brain (B) and in vivo human brain (C).

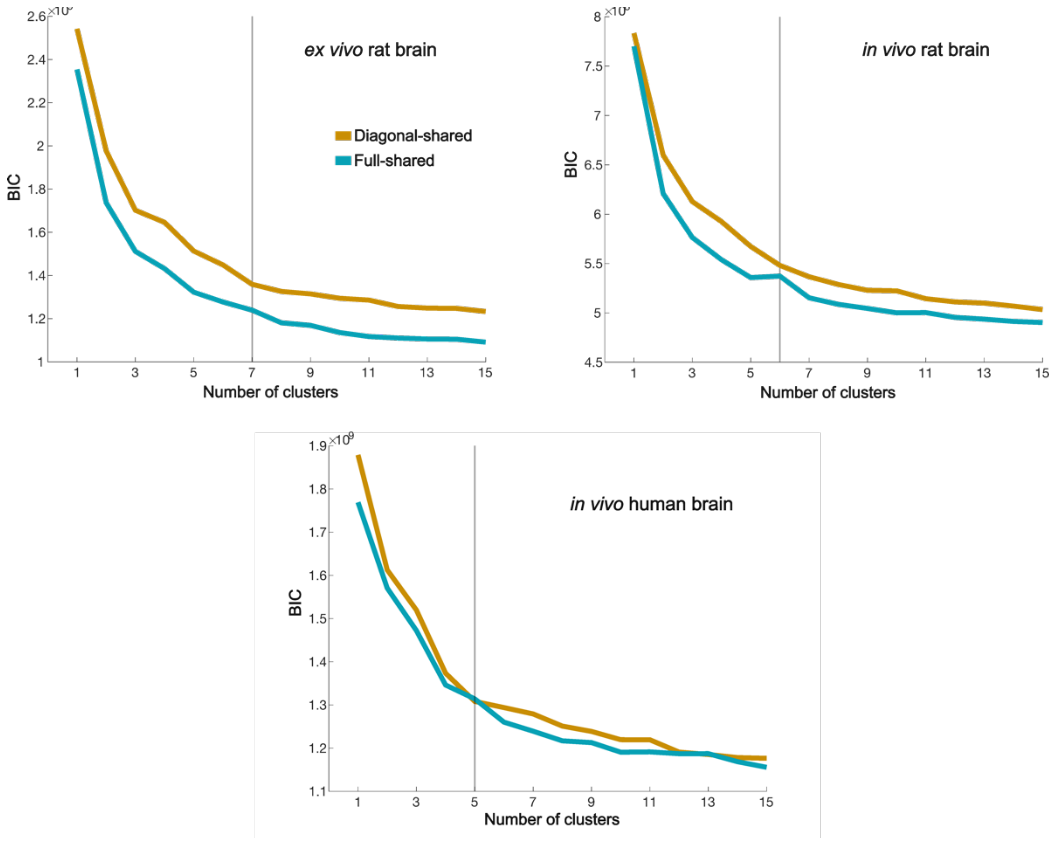


**Supplementary Figure 2.** Schwarz's Bayesian Information Criterion (BIC) from the clusterization of the $D_{\mathrm{iso}}$, $D_{\Delta}$, $R_1$, $R_2$ and $\Delta_{\omega/2\pi}D_{\mathrm{iso}}$ space of *ex vivo* rat brain, $D_{\mathrm{iso}}$, $D_{\Delta}$, $R_1$, $R_2$ and $\Delta_{\omega/2\pi}D_{\mathrm{iso}}$ space of *in vivo* rat brain, and $D_{\mathrm{iso}}$, $D_{\Delta}$, $R_1$, and $R_2$ space of *in vivo* human brain.

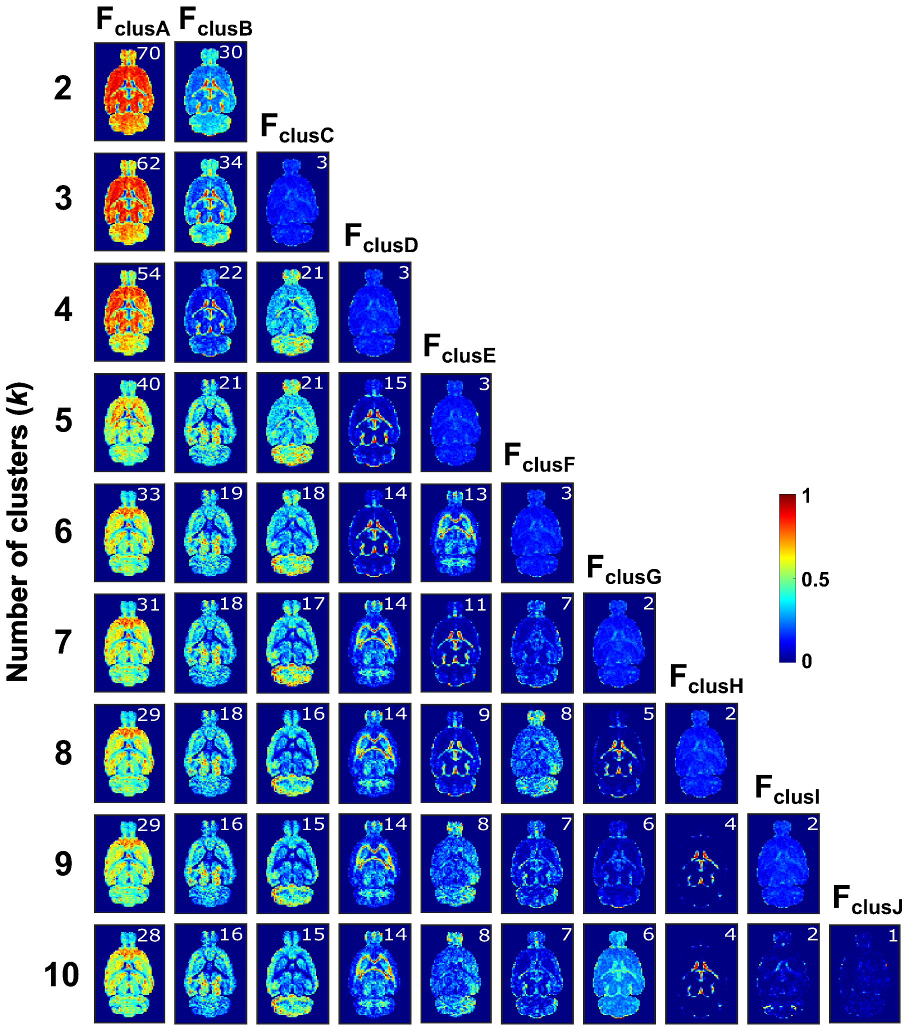


**Supplementary Figure 3.** Normalized cluster frequency of $k = 2$-10 from the full distribution space ($D_{\mathrm{iso}}$, $D_{\Delta}$, $R_1$, $R_2$ and $\Delta_{\omega/2\pi}D_{\mathrm{iso}}$) of *in vivo* rat brain data. Rows show the number of clusters and columns the normalized cluster frequency. The clusters are ordered by decreasing percentage of weight belonging to the cluster from left to right. The color scale shows the per-voxel weight belonging to each cluster.

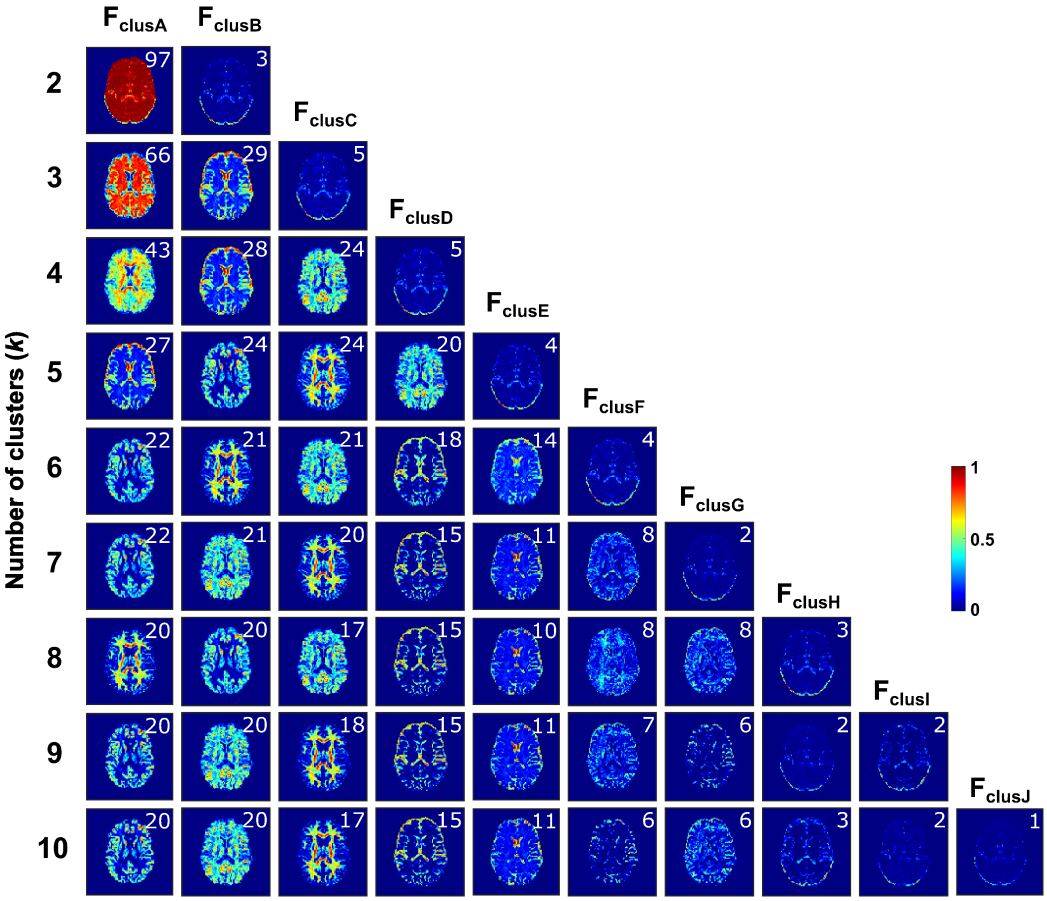


**Supplementary Figure 4.** Normalized cluster frequency of $k = 2$-10 from the $D_{iso}$, $D_{\Delta}$, $R_1$, and $R_2$ distribution space of *in vivo* human data. Rows show the number of clusters and columns the normalized cluster frequency. The clusters are ordered by decreasing percentage of weight belonging to the cluster from left to right. The color scale shows the per-voxel weight belonging to each cluster.
